# Higher-order thalamic circuits channel parallel streams of visual information in mice

**DOI:** 10.1101/395244

**Authors:** Corbett Bennett, Samuel D. Gale, Marina E. Garrett, Melissa L. Newton, Edward M. Callaway, Gabe J. Murphy, Shawn R. Olsen

## Abstract

Higher-order thalamic nuclei, such as the visual pulvinar, play essential roles in cortical function by connecting functionally-related cortical and subcortical brain regions. Yet a coherent framework describing pulvinar function remains elusive due to its anatomical complexity and involvement in diverse cognitive processes. Here we combined large-scale anatomical circuit mapping with high-density electrophysiological recordings to dissect a homolog of pulvinar in mice, the lateral posterior nucleus (LP). We define three broad LP subregions based on correspondence between input/output connectivity and functional properties. These subregions form corticothalamic loops biased towards ventral or dorsal stream cortical areas and contain separate representations of visual space. To reveal which input sources drive LP activity, we silenced visual cortex or superior colliculus and found they drive visual tuning properties in separate LP subregions. Thus, by specifying the driving input sources, functional properties, and downstream targets of LP circuits, our data provide a roadmap for understanding the mechanisms of higher-order thalamic function in vision.

## Introduction

Higher-order thalamus is critically involved in cortical function, both as a route by which cortical areas communicate and a relay of subcortical input to cortex. The pulvinar, the higher-order thalamic nucleus in the visual system, is reciprocally connected with multiple visual and frontal cortical areas and therefore mediates an indirect pathway for cortico-cortical communication through cortico-thalamo-cortical (“transthalamic”) circuits (Sherman, 2016). The pulvinar also receives a dense projection from the superficial (retino-recipient) layers of the superior colliculus (SCs) which interacts with transthalamic circuits, potentially forming a secondary route of information flow from retina to cortex that runs parallel to the retino-geniculate pathway. The relative contributions of cortical and SCs input in driving pulvinar visual responses and the information these pathways carry remain uncertain.

The lateral posterior thalamic nucleus (LP) in rodents is thought to be homologous to the primate visual pulvinar (Baldwin et al., 2017). Like the pulvinar, mouse LP is reciprocally connected with all visual cortical areas and receives a strong projection from SCs (Oh et al., 2014; Tohmi et al., 2014; Zhou et al., 2017). Previous studies divided LP into subregions based on cytoarchitecture and immunocytochemical markers (Nakamura et al., 2015; Takahashi, 1985; Zhou et al., 2017), but it is unclear whether these subregions reflect functionally distinct domains of LP. Conversely, functional studies have shown that LP neurons encode a variety of sensory and motor signals and convey these to the visual cortex (Allen et al., 2016; Durand et al., 2016; Roth et al., 2016), but these studies focused on limited portions of LP and did not relate functional differences to LP anatomy. Moreover, none of these studies tested the causal role of cortical or SC input in driving LP visual responses.

Here, we performed comprehensive anatomical input/output mapping and high-density electrophysiological recordings across the full extent of LP to elucidate the relationship between structural and functional organization in the mouse corticothalamic visual system. We find that LP is not a homogenous structure—rather, we identify three subregions with different patterns of connectivity with visual cortical areas, frontal cortex, and SC. To test whether these regions differentially depend on the SC or retino-geniculate-V1 pathways, we silenced either SC or primary visual cortex (V1). We find that SC input drives visual responses to object motion in posterior LP. Interestingly, unlike in primates (Kaas and Lyon, 2007), this information is routed primarily to ventral stream rather than dorsal stream visual cortical areas. In contrast, visual responses in anterior LP require V1 but not SC input and are primarily routed to dorsal stream visual cortical areas. Together these data show a tight correspondence between anatomical, functional, and perturbational maps of LP, and reveal that SC is the main driver of activity in a thalamo-cortical circuit linking LP to primarily ventral stream visual areas.

## Results

### Anatomical input mapping divides LP into three broad subregions

LP activity is shaped by input from diverse cortical and subcortical structures. Functionally-relevant subregions of LP might therefore be defined by distinct patterns of input connections. To generate detailed maps of LP input connectivity, we quantified axon labelling in LP resulting from anterograde viral tracer injection into SCs, eight visual cortical areas, and two frontal cortical areas (Fig. 1A; all injections taken from Allen Mouse Brain Connectivity Database). Axonal projections were mapped to a standardized anatomical coordinate system for comparison (the Allen Institute Common Coordinate Framework, CCF), yielding an “input volume” for each source area representing which LP voxels are targeted by axons from that area.

**Figure 1.**
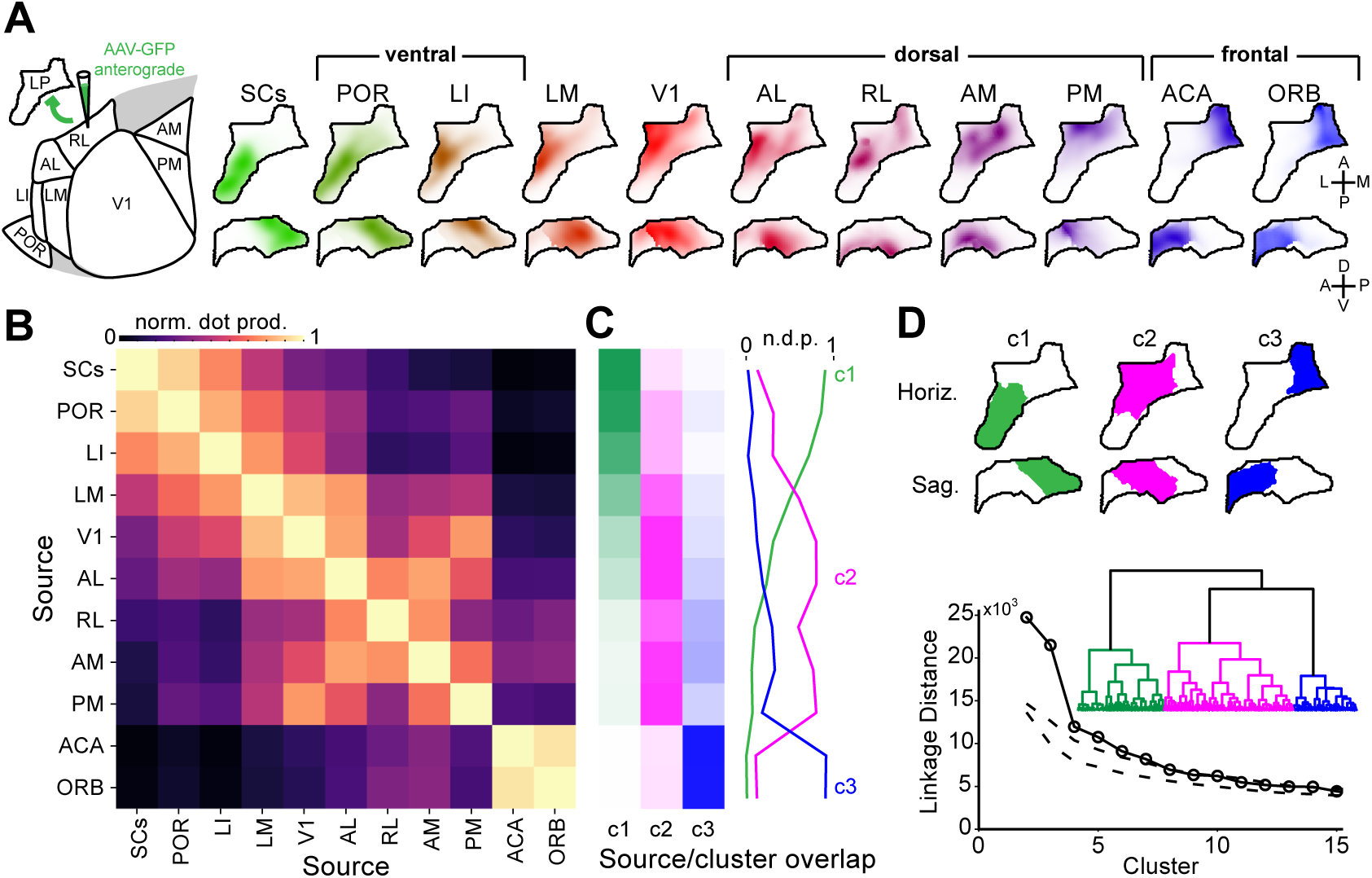
Input connectivity reveals three broad LP subregions. **(A)** Left: schematic of input mapping experiments showing visual cortical areas in the horizontal plane. Right: location of anterogradely labeled axons in LP from various cortical and subcortical input sources. Input volumes are shown as horizontal (top row) and sagittal projections (bottom row) and represent average fluorescence across multiple tracer injection experiments registered in the Allen Institute Common Coordinate Framework (see Methods). **(B)** Overlap of inputs to LP from different sources (normalized voxel-wise dot product). **(C)** Overlap of input from each source with clusters of LP voxels based on all inputs (hierarchical clustering using Ward’s linkage criterion). **(D)** Top: projections of LP voxels belonging to the first three clusters. Bottom: Dendrogram (inset) shows linkage distance of LP voxels based on anatomical inputs. Linkage distance for the first 15 clusters are compared to clusters formed from random shuffling of data across voxels for each input source (dashed lines; 1–99% confidence interval). Abbreviations: superficial superior colliculus (SCs), postrhinal area (POR), laterointermediate area (LI), lateromedial area (LM), primary visual cortex (V1), anterolateral area (AL) rostrolateral area (RL), anteromedial area (AM), posteromedial area (PM), anterior cingulate (ACA), orbital cortex (ORB).

Visual inspection of LP input volumes revealed two general principles of LP organization. First, SCs axons projected densely to the posterior half of LP but were sparse in anterior regions, whereas input from frontal areas (ACA and ORB) was confined almost exclusively to medial LP. Second, input from visual cortical areas were overlapping but organized such that axons from ventral stream areas tended to aggregate in posterior and dorsal LP with SCs axons, while axons from dorsal stream areas were biased towards anterior and ventral LP along with V1 axons (Fig. 1A).

To quantify the degree to which axons from different source regions overlap in LP, we computed the normalized dot product between each input volume (Fig. 1B). The resulting similarity matrix confirmed our qualitative observations and suggested that specific groups of input sources target different regions of LP. To further quantify these relationships, we clustered LP voxels based on their input (Fig. 1C,D). Hierarchical or k-means methods produced nearly identical clusters (>96% of voxels were the same for each cluster). This analysis revealed three statistically significant clusters that differed in their predominant sources of input: (1) posterior/dorsal LP (pLP; cluster 1) receives input primarily from SCs and ventral-stream visual cortical areas (LI and POR); (2) anterior/ventral LP (aLP; cluster 2) receives input primarily from V1 and dorsal-stream visual cortical areas (AL, RL, AM, PM); and (3) medial LP (mLP; cluster 3) receives input from frontal cortical areas. Importantly, projections from any given cortical area were biased towards but not exclusive to one of these three clusters; the clusters are therefore defined by dissimilar but overlapping patterns of input.

Similar patterns of axonal labeling were observed in LP when we limited our analysis to smaller visual cortical injections guided by intrinsic signal imaging, or when fluorophore expression was restricted to synaptic terminals (rather than fibers of passage) or to layer 5 or 6 cortical neurons (in Rbp4-KL100 or Nstr1-GN220 Cre mice, respectively; Fig. S1C-J). Input from layer 5 and 6 neurons from a given area overlapped more with each other than layer 5 and 6 input from separate areas (Fig. S1K).

### LP output mapping reveals reciprocal and relay transthalamic pathways

What are the projection targets of these LP subregions? We defined LP output volumes by measuring fluorescence from LP cells retrogradely labelled by rabies injections into ten target regions, including the amygdala and nine of the cortical areas from the input analysis (Fig. 2A; S2A-H). These output volumes were mapped to the CCF for comparison with input volumes.

**Figure 2.**
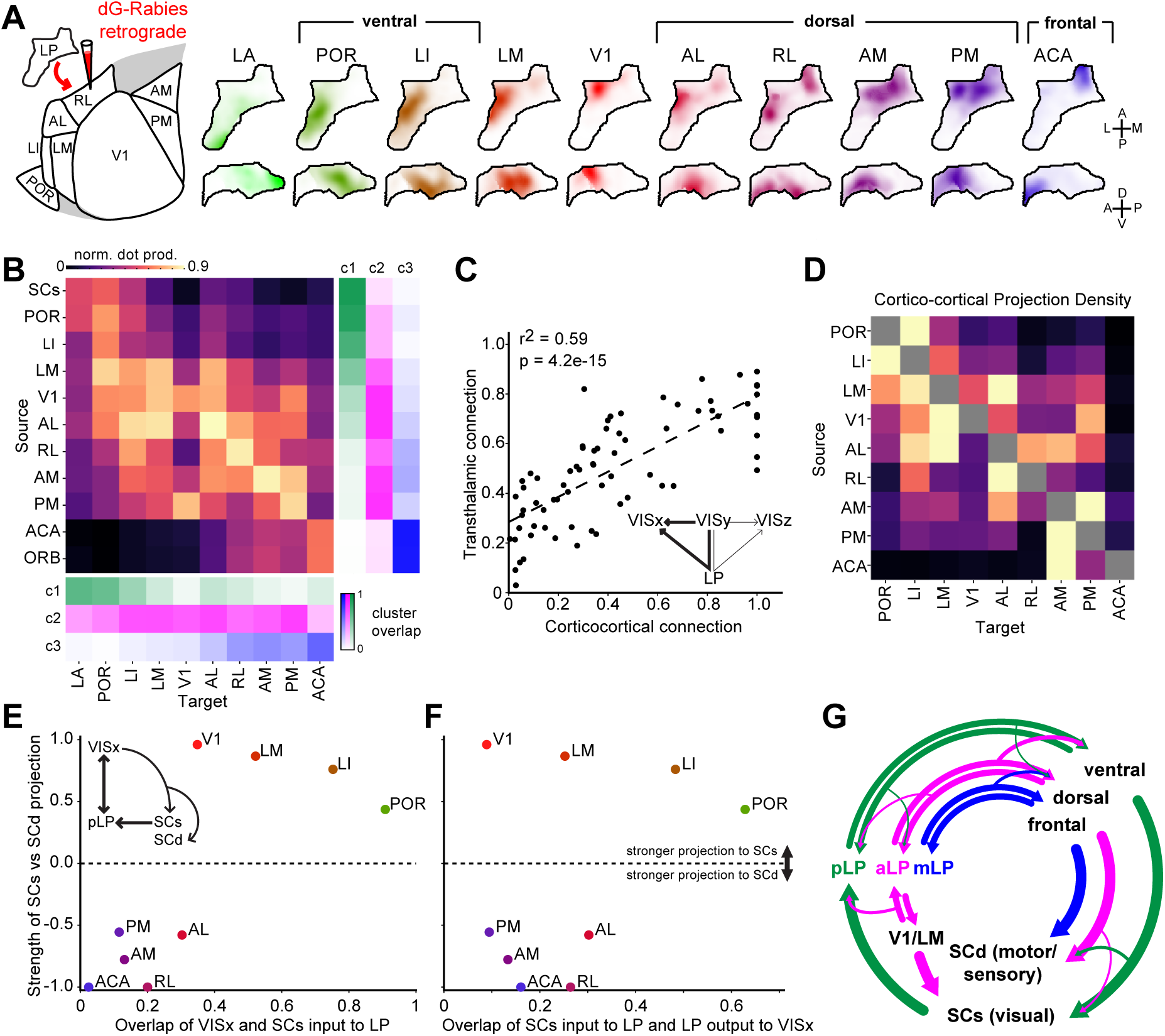
LP input/ouput mapping reveals reciprocal and relay transthalamic pathways. **(A)** Location of retrogradely labeled cells in LP following rabies injections in various output targets. Output volumes are shown as horizontal and sagittal projections as for input volumes shown in Fig. 1A. **(B)** Overlap (normalized dot product) in LP of input and output volumes for each source-target region pair. Right: overlap of input volumes and clusters from Fig. 1C are shown again here for reference. Below: overlap of output volumes and the clusters from Fig. 1. **(C,D)** Comparison of direct cortico-cortical and indirect cortico-LP-cortical (transthalamic) connectivity. The density of axons directly connecting visual cortical areas (panel D; cortico-cortical connections, row-normalized) are compared in (C) to the overlap of input from and output to the same source-target pairs in LP (panel B; putative transthalamic connections). **(E)** Relative strength of cortical projections to SCs or deep superior colliculus (SCd) as a function of overlap between cortical and SCs LP projections for nine cortical areas (x-axis taken from top row of Fig. 1B). **(F)** Same as E but for overlap of SCs input and LP output to the same cortical areas (x-axis taken from top row of B). Values on y-axis for E,F are the difference of the projection density to SCs and SCd divided by their sum (Methods). **(G)** Diagram summarizing connectivity between LP, cortex, and SC.

To quantify LP input/output connectivity, we compared the overlap of anterogradely-labeled axons and retrogradely-labeled cell bodies from each input source and output target (Fig. 2B). This analysis revealed putative transthalamic pathways linking visual areas by indirect projections through LP. We found that most of these pathways are reciprocal: if an LP voxel receives input from a region, it tends to project back to that region (evident from high values along the diagonal of Fig. 2B). However, we also found evidence for relay pathways linking distinct areas (evident from values off the diagonal). For example, mLP voxels (cluster 3) receive input almost exclusively from frontal cortical regions (ACA and ORB) and project back to these regions as well as to dorsal-stream visual cortical areas (PM, AM, RL, AL). Similarly, aLP voxels (cluster 2) receive most of their input from dorsal-stream visual cortical areas but project both back to these regions and to ventral-stream visual cortical areas (LI and POR). Finally, pLP voxels (cluster 1) project to the amygdala (LA) in addition to reciprocal connections with LI and POR, but do not project back to SCs.

A complementary analysis using maps of LP output defined by anterograde tracer injections in different portions of LP yielded similar results (Fig. S2I-K). This analysis also revealed which layers LP axons target in each cortical area. We found that injections into anterior and posterior LP produced similar patterns of axons in their respective cortical targets (Fig. S2L). Interestingly, however, the layer distribution of LP axons differed across cortical areas. To explain this variability, we ordered areas by their position in the cortical hierarchy, according to a recently published metric (Harris et al., 2018). We found that areas lower in the cortical hierarchy received a larger proportion of LP axons in layer 1, while areas higher in the hierarchy received a larger proportion input in layer 4 (Fig. S2M).

Cortical areas are connected directly through cortico-cortical projections and indirectly through higher-order thalamic regions such as LP (transthalamic pathways). To determine whether the same or different visual cortical areas are connected via cortico-cortical and transthalamic pathways (Fig. 2C, inset), we compared: (1) the density of axons from each cortical area to all the other cortical areas (cortico-cortical projections; Fig. 2D), with (2) the overlap of axons in LP from each cortical area (input volumes) and the cell bodies in LP (output volumes) projecting to each of those areas (transthalamic connections; Fig. 2B). There is a significant correlation between these measures of cortical connectivity (Fig. 2C), suggesting that transthalamic connections of visual cortical areas through LP largely mirror direct cortico-cortical pathways. Additionally, the greater overlap evident in the transthalamic connectivity matrix indicates that LP may also supplement direct cortico-cortical connectivity by providing additional routes of communication between cortical areas.

Finally, LP is known to project to the dorsal striatum; however, little is known about the topography of this projection. Analysis of injections into LP from the Allen Connectivity Database revealed a coarse topography of LP output to the dorsal striatum, with more posterior regions of LP targeting posterior-lateral striatum and more anterior regions in LP targeting anterior-medial striatum (Fig. S2L).

### Ventral stream areas participate in cortico-collicular-thalamic loops

All visual cortical areas in mice project to SCs as well as to deeper SC layers (SCd). Cortical projections exhibit stream specific biases to SCs (ventral stream areas) or SCd (dorsal stream areas) (Wang and Burkhalter, 2013), and may provide a route by which some visual cortical areas can indirectly influence their input from LP (Fig. 2E inset). To identify potential loops connecting LP, cortex, and SC, we compared: (1) the relative strength of input from each cortical area to SCs and SCd, with (2) the overlap of SCs input to LP with each cortical area’s input to or output from LP (the top rows of Fig 1B and 2B, respectively). We found that the ventral stream areas (LI and POR) that project strongly to SCs are also strongly interconnected with the SCs-recipient portion of LP (Fig 2E,F). In contrast, dorsal stream and frontal areas that project strongly to SCd are less connected with the SCs-recipient portion of LP. The geniculo-recipient cortical areas (V1 and LM) project strongly to SCs like ventral stream areas, but similar to dorsal stream areas are relatively weakly connected with SCs-recipient LP. Thus, ventral stream areas are the main cortical component of a trisynaptic loop connecting pLP, cortex, and SCs (Fig. 2G).

Conversely, we find evidence for only a weak projection to LP from SCd. Allen Connectivity Database injections that more strongly overlapped SCd than SCs labeled relatively sparse axons in pLP (probably due to the small amount of virus in SCs) and mLP (Fig. S1L), suggesting that SCd projects to mLP and that anterior LP does not receive input from SCs or SCd. However, the SCd projection to mLP is much sparser than the SCs projection to pLP.

### LP contains at least two maps of visual space corresponding to posterior (SC-recipient) and anterior (V1-recipient) subregions

If the LP subregions we defined anatomically are functionally distinct, they may, like visual cortical areas, have separate maps of visual space. Indeed, SC and V1 contain full representations of visual space but project to largely non-overlapping parts of LP (Fig. 1A,B). We used the retinotopic organization of SC and V1 and the topography of their axons in LP to predict the representation of visual elevation and azimuth in LP (Fig. 3A-F, S3). The predicted elevation map showed separate spatial gradients of visual elevation representation in LP that converge at the boundary of SC and V1 input to LP. Thus, anatomical connectivity predicts that LP contains at least two distinct representations of visual elevation.

**Figure 3.**
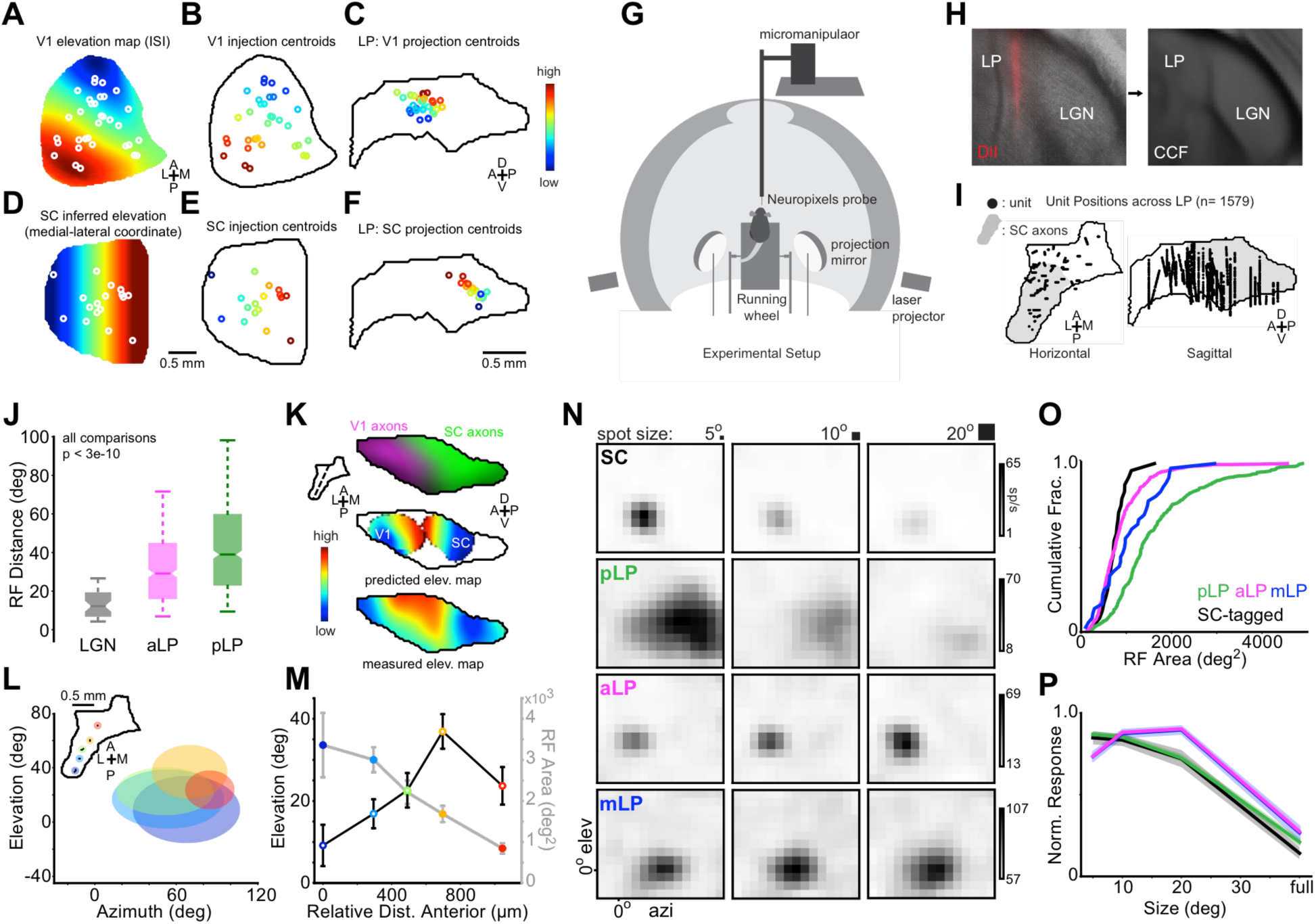
Posterior and anterior LP have separate maps of visual space and distinct receptive field properties. **(A)** Mean ISI elevation map for V1 with location of V1 injections from the Allen Connectivity Database superimposed (white circles indicate injection centroids). **(B)** V1 injections colored by assigned elevation according to map in (A). **(C)** V1 projection centroids in LP (sagittal plane) for injections in (B) colored by assigned elevation. These centroids were smoothed to create the V1 predicted elevation map in (K). **(D-F)** Same as (A-C) for SC injections. Elevation in SC is inferred from the medial-lateral coordinate (Methods). **(G)** Diagram of experimental setup for visual stimulation and neural recording. **(H)** DiI labeling of probe tract recovered from post-hoc histology and registered to the CCF. **(I)** Recording locations for all LP neurons displayed on horizontal and sagittal projections of LP. Gray region denotes SC-recipient LP. **(J)** Receptive field distance for pairs of cells separated by 20 microns or less in dLGN (gray, n=48 pairs), aLP (magenta, n=350 pairs) or pLP (green, n=554 pairs). Only cells from the same probe insertion were compared. Box edges indicate first and third quartiles. Notch indicates 95% CI for median (band). Whiskers denote 5th and 95th percentiles. Wilcoxon rank-sum test. **(K)** Top: LP slice showing SC (green) and V1 (magenta) input to LP. Plane of slice indicated by dotted line in inset. Middle: Predicted LP elevation map based on anatomical V1 input (left) or SC input (right). Bottom: Composite elevation map for all LP cells. (**L)** Data from experiment in which five insertions were made serially in one mouse. Recording locations for each insertion are shown in the inset. Ellipses are centered at the mean RF center for each insertion (color-coded to match inset). Ellipse shape reflects mean RF shape for neurons at each location. **(M)** Mean receptive field area (closed circles) and elevation (open circles) for each recording location in (L). Colors as in (L). Error bars represent standard error. **(N)** Off receptive fields for example SC (optotagged), pLP, aLP and mLP neurons (rows). Receptive fields were mapped with sparse noise consisting of 5, 10, and 20 degree squares (columns). **(O)** Cumulative distribution of receptive field area and **(P)** mean size tuning for SC (black), pLP (green), aLP (magenta) and mLP (blue) neurons. Shaded regions in (P) denote standard error.

To test this prediction, we used Neuropixels high-density electrode probes (Jun et al., 2017) and a sparse noise stimulus to measure the spatial receptive field location of neurons over the full extent of LP in awake mice (Fig. 3G). The location of each probe in LP was marked with DiI, and every recorded cell was assigned a position in the CCF (Fig. 3H,I; 1579 neurons, Methods). The dispersion of receptive field locations of nearby neurons was substantially higher in LP than the dorsal lateral geniculate nucleus (dLGN; Fig. 3J). Nonetheless, averaging the receptive field locations of nearby LP neurons revealed a smooth map of elevation that, like the elevation map predicted by anatomy, reversed its gradient near the center of LP where SC and V1 axons converge (Fig. 3K). We confirmed our findings from population data by serially recording responses to sparse noise in multiple locations across LP in the same mouse (Fig. 3L). As we moved the probe to more anterior locations, the receptive field elevation gradient reversed (Fig. 3M). This data strongly suggests that pLP and aLP are functionally distinct subregions containing separate maps of visual space.

Similar methods were used to construct maps of azimuth representation in LP. Both the topography of axons and the measured receptive fields showed a gradient of azimuth representation in the region of LP receiving input from V1 but no clear azimuth map in the region of LP receiving SC input (Fig. S3).

Mapping receptive field organization in pLP is potentially complicated by the fact that this region contains adjacent zones receiving unilateral or bilateral SC input (Zhou et al., 2017). In other species, SC axons are topographically organized in the part of LP receiving unilateral input but more diffuse in the part receiving bilateral input (Baldwin et al., 2011, 2013; Chomsung et al., 2008). We found that SC axons are topographically organized for elevation but not azimuth in both the unilateral and bilateral SC-recipient portions of LP (Fig. S4), consistent with data from rats (Takahashi, 1985). However, this organization was more precise in unilateral SC-recipient LP (Fig. S4H).

We find evidence for three LP subregions with distinct representations of visual space. What rules govern the representation of each visual cortical area within these subregions? Visual cortical areas in mice often contain incomplete retinotopic maps biased to a particular region of visual space (Garrett et al., 2014; Zhuang et al., 2017). These biases might be reflected in the organization of cortical connections with LP. We computed the elevation bias for the portion of LP receiving input from, or projecting to, each cortical area by weighting the LP elevation map by the anatomical input and output volumes shown in Figures 1 and 2. We found a significant correlation between these elevations and the mean elevation measured for each cortical area via intrinsic imaging (Fig. S5). This result indicates that the LP input and output maps associated with a given visual cortical area are in part predicted by that area’s retinotopic bias.

### Mapping functional properties across LP with high-density electrophysiology

By registering our LP recordings to the CCF, we were able to map visual response properties across LP and compare these functional maps to the anatomically-defined subregions we describe above. We compared visual responses in pLP (the subregion that receives dense input from SC), mLP (the subregion that receives dense input from ACA), and aLP (sparse if any input from SC or ACA). In addition, we compared visual responses in LP to putative LP-projecting SC neurons recorded in Nstr1-GN209 Cre x Ai32 mice. In these mice, the SC cell type that projects to LP, but not other SC cell types, expresses channelrhodopsin-2 (ChR2) and responds with sustained spiking to long flashes of blue light (Gale and Murphy, 2016). There is minimal if any connections between these LP-projecting cells and other SC cell types (Gale and Murphy, 2018). Consistent with previous results, visual responses of optotagged SC neurons differed substantially from those of non-optotagged SC neurons recorded in the same mice (Fig. S6). By characterizing visual response properties of neurons in the three LP subregions and identified LP-projecting SC neurons, we are able not only to map functional differences across LP, but also to elucidate how visual information in SC is transformed in LP before it is conveyed to visual cortex.

### Receptive field size and size tuning differ across LP subregions

Reponses to sparse noise consisting of black and white squares of varying size revealed sub-region specific differences in the size and shape of spatial receptive fields and size tuning in LP. Receptive fields in pLP were horizontally elongated and significantly larger than aLP and mLP receptive fields (Fig. 3N,O; Table S3; median aspect ratio: 1.30 pLP, 1.15 aLP, 1.05 mLP, 1.07 SC). Despite their large spatial receptive fields, pLP neurons, like optotagged SC neurons, responded most strongly on average to the smallest square size presented (5°; Fig. 3N,P). These neurons thus respond best to small stimuli presented at any location within a relatively large region of space. In contrast, aLP and mLP neurons responded most strongly to the largest square size (20°). These results are consistent with functional differences, in addition to anatomical differences, between LP subregions.

### Responses to object and background motion differ in LP subregions

LP-projecting cells in SC respond more strongly to small, slowly-moving objects in the visual field than to full-field background motion (Gale and Murphy, 2016). To determine whether LP neurons respond differentially to object or global motion, we recorded responses to a stimulus consisting of a moving random checkerboard background (full field) and a small (10°), differentially-moving “patch” of random checkerboard pattern. Because the patch is only visible when its speed and/or direction of motion differs from that of the background, this stimulus is well-suited to probing responses to object versus global motion (Frost et al., 1988). Responses to a matrix of 7 patch and 7 background velocities were recorded (Fig. 4A-D). To visualize and compare responses across cells, we compiled an N cells (1135) by 49 condition (patch and background velocity combinations) population response matrix (Fig. 4E). Hierarchical clustering on this matrix revealed three main clusters among which cells from SC and the three LP subregions were differentially distributed (Fig. 4E, right).

**Figure 4.**
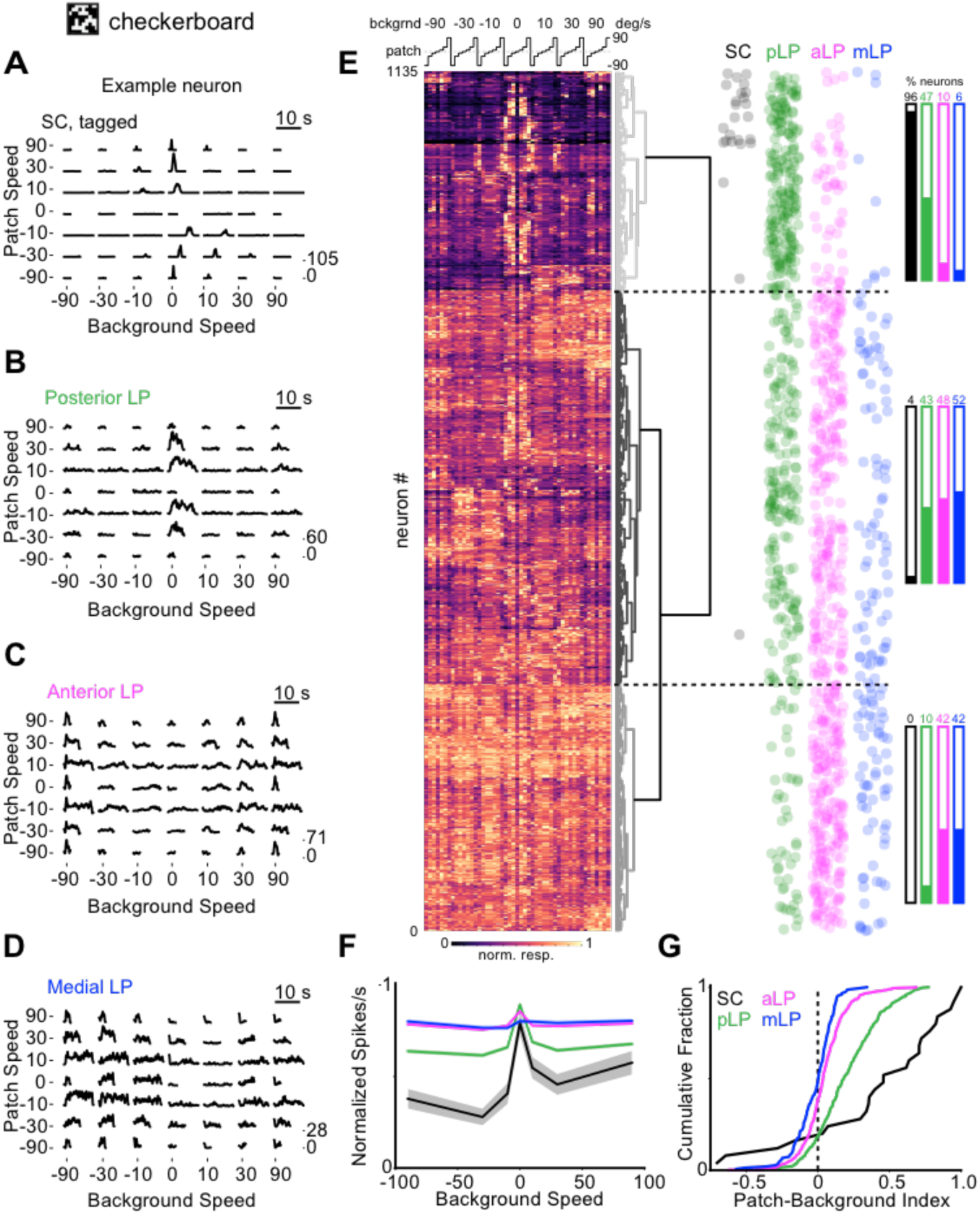
LP neurons differ in their response to object and background motion. **(A)** Spike density functions of an example optotagged SC neuron (left) and mean population response (right) to random checkerboard background (full field) and patches (10°) moving relative to each other at various speeds (positive speeds are nasal to temporal). Checkerboard squares are 1°. Patch speed 0°/s trials are background motion only. Background 0°/s trials consist of patches moving over a stationary random checkerboard background. Since the texture of patches and background are indistinguishable, patches moving with the same speed and direction as background are invisible (equivalent to patch speed 0°/s trials). **(B,C,D)** Same as (A) for pLP, aLP, mLP neurons. **(E)** Left: heat map showing the normalized response of all SC and LP neurons to the checkerboard stimulus. Each row represents one cell. The matrix of 7 patch and 7 background speeds shown in (A-D) are linearized to a 49 element vector as shown above the heat map. Hierarchical clustering was used to order the rows according to the linkage distance between cells (represented by the dendrogram to the right of the heat map). Right: dot plots showing the position (row) of each cell from SC or an LP subregion along the heat map. The horizontal location of the dots were jittered randomly to reduce overlap. Bar plots indicate the percentage of cells from each region belonging to the three main clusters in the heat map data. **(F)** Population tuning curves for background speed (normalized max response down columns of checkerboard response matrix) for SC (black), pLP (green), aLP (magenta) and mLP (blue). **(G)** Cumulative distributions of patch-background index values for SC, pLP, aLP and mLP in response to the checkerboard stimulus. Patch-background index is the difference between the maximum responses to patch (background speed 0°/s) and background (patch speed 0°/s) motion divided by their sum.

LP-projecting SC neurons almost exclusively belonged to a cluster of cells that responded most strongly to the patch moving over a stationary background (top cluster with relatively high values in the middle columns of Fig. 4E). In some cases, these neurons also responded moderately when the patch and background moved in opposite directions and/or with a large difference in speed. Neurons in pLP were similar to SC, but more likely to be modulated by background motion, and were split between the patch selective cluster (top) and the middle cluster, in which many cells exhibited preferences for the direction and/or speed of patch and background motion. By contrast, cells in aLP and mLP almost exclusively belonged to this middle cluster or the bottom cluster in which cells were visually responsive but not selective.

Responses to patch versus background motion were summarized by two metrics: (1) plotting response size as a function of background motion (Fig. 4F), and (2) calculating a “patch-background index” comparing maximum responses to patch or background motion alone (Fig. 4G). Consistent with the clustering shown in Fig. 4E, SC and pLP neurons, on average, responded maximally when the patch moved over a stationary background (background speed 0 °/s; Fig. 4F) and, unlike aLP and mLP neurons, exhibited a strong bias for positive patch-background index values (indicating preference for patch motion; Fig. 4G; Table S3). Overall, this data further implies that pLP is functionally distinct from aLP and mLP, and, along with size tuning data (Fig. 3N,P), is consistent with the possibility that input from SC plays an important role in shaping visual responses of pLP neurons.

Responses to full-field drifting gratings revealed additional differences in motion sensitivity between neurons in LP subregions. Neurons in aLP and mLP typically preferred faster speeds and higher temporal frequencies than pLP neurons (Fig. S7). Direction-selective neurons in pLP tended to prefer motion towards the upper-temporal visual field similar to the bias found in SC (Ahmadlou and Heimel, 2015; Drager and Hubel, 1975; Gale and Murphy, 2014) (Fig. S7G), while neurons in aLP and mLP tended to prefer motion towards the nasal visual field (Fig. S7H-J).

### Posterior LP neurons respond more strongly to looming stimuli

The portion of LP that receives input from SC and projects to the amygdala has been implicated in mediating a freezing response to looming stimuli (Shang et al., 2018; Wei et al., 2015). To characterize responses of LP neurons to looming stimuli, we presented expanding spots that increased in angular size at a rate that simulates an object approaching at constant speed; the rate of expansion is a nonlinear function of time-to-collision and the object’s size/speed ratio (Gabbiani et al., 1999) (Fig. 5A). SC and pLP neurons responded more strongly to looming stimuli than aLP or mLP neurons (Fig. 5C; S8A). Most cells that responded to looming stimuli also responded to the checkerboard stimulus (23/23 SC, 110/113 pLP, 11/13 aLP, 22/26 mLP). In many SC and pLP neurons, the time of the peak response to looming stimuli was linearly related to the object size/speed ratio such that the peak response occurred at a constant angular spot size (Fig. 5B). Similar responses to looming stimuli are observed in other animals and are referred to as η cell responses (de Vries and Clandinin, 2012; Gabbiani et al., 1999; Liu et al., 2011; Sun and Frost, 1998). Other types of responses (e.g. ρ and τ cells) were not clearly observed in our experiments (Fig. S8B). We define η cells as those with correlations greater than 0.9 between the time of peak response and the object size/speed ratio. Cells defined as η cells were more common in pLP than other parts of LP (64% of LP-projecting SC, 25% of pLP, and 5% of aLP and mLP neurons; Fig. 5D,E). Within pLP, other properties of η cells also differed from non-η cells. In response to the checkerboard stimulus, pLP η cells exhibited a stronger preference for object (patch) motion compared to background motion than non-η cells (Fig. 5F). Thus, a population of pLP neurons may convey information about looming stimuli, and object motion in general, to the amygdala and ventral stream visual cortical areas.

**Figure 5.**
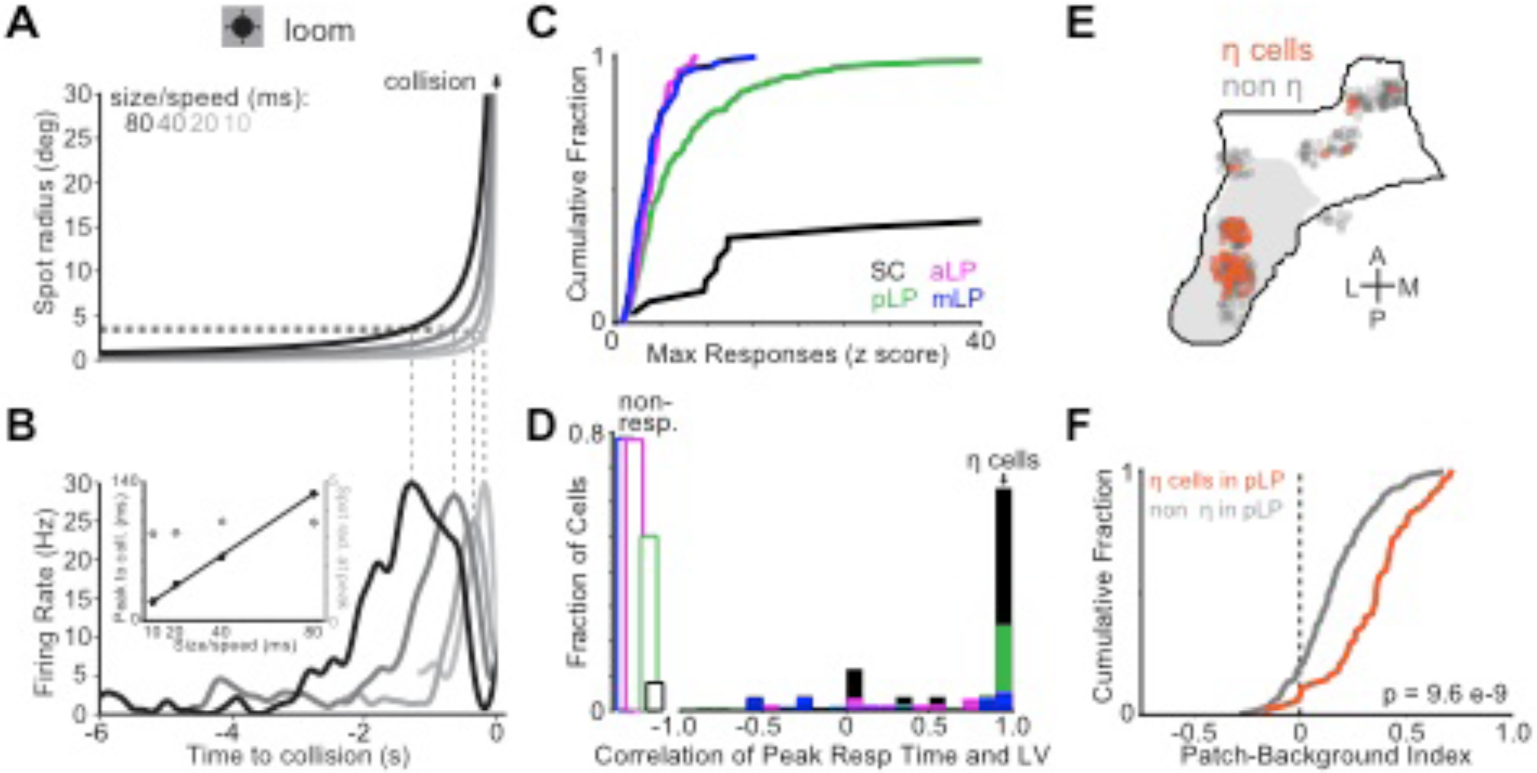
SC and posterior LP neurons respond more strongly to looming stimuli. **(A)** Trajectory of spot radius for looming stimuli at four size-to-speed ratios. **(B)** Firing rate of example neuron in posterior LP to looming stimuli depicted in (A). Dotted lines relate spot radius to time of peak firing rate. Inset: time of peak firing rate relative to collision plotted against size to speed ratio for example neuron in (B) (filled black circles; left axis) and spot radius at peak firing rate plotted against size to speed ratio for same neuron (open gray circles; right axis). **(C)** Cumulative distribution of max loom response (z score) across all conditions for neurons in SC (black), pLP (green), aLP (magenta), and mLP (blue). Note that, due to their low spontaneous firing rates, many SC neurons had z-scores greater than 40. **(D)** Histogram of correlation between peak response time and size to speed ratio for cells in SC and LP subregions (colors as in (C)). Cells with a correlation value greater than 0.9 were classified as η-type. Open rectangles on left represent fraction of cells that were not responsive to looming stimuli. **(E)** Location η cells shown in a horizontal projection of LP. Gray region denotes SC-recipient LP. **(F)** Cumulative distribution of checkerboard patch-background index values for η (orange, n=55 cells) and non-η (gray, n=150 cells) neurons in posterior LP. Wilcoxon rank-sum test.

### Clustering LP neurons based on visual response properties alone independently segregates posterior and anterior neurons

Our results demonstrate that distinct patterns of anatomical connections segregate LP into subregions that contain neurons with different functional properties. Is the inverse also true: do functional properties predict the spatial organization of neurons in LP? Hierarchical clustering of LP neurons based on visual response properties (size tuning, RF area, and patch-background index) produced two statistically significant clusters (Fig. 6A). Compared to neurons in cluster 1, neurons in cluster 2 had larger receptive fields, stronger surround suppression, and stronger preference for object motion compared to background motion (Fig. 6C-E). These properties are similar to those of neurons located in pLP. Indeed, 52% of pLP neurons were assigned to cluster 2, while only 7% of aLP neurons and 3% of mLP neurons belonged to this cluster (Fig. 6B). Thus, functional properties independently predict spatial segregation of neurons in LP.

**Figure 6.**
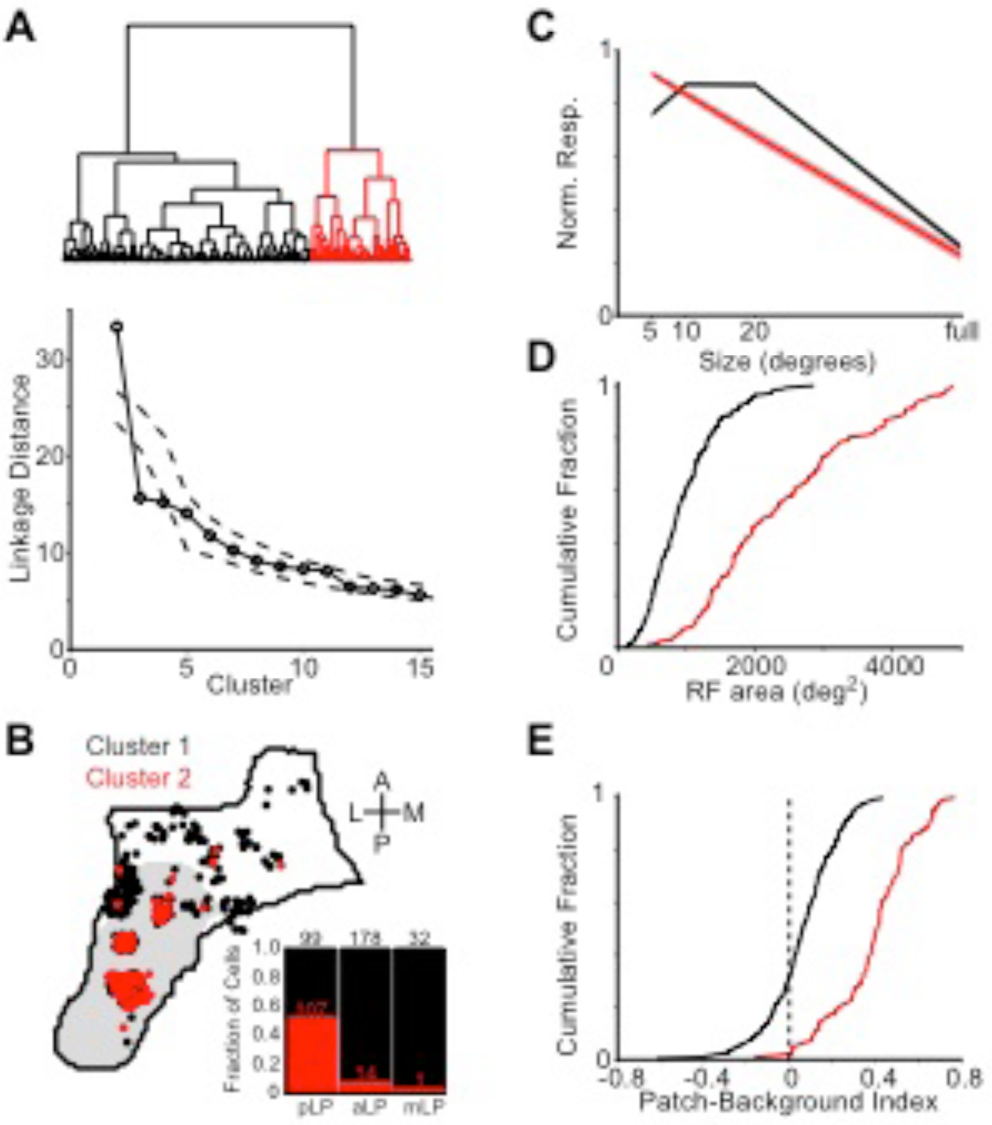
Visual response properties reveal functional-anatomical segregation in LP. **(A)** Top: dendrogram representing hierarchical clustering (Ward’s linkage criterion) of LP neurons based on visual response properties. Bottom: linkage distance for the first 15 clusters compared to clusters formed from the same data randomly shuffled across neurons for each visual response parameter (dashed lines are 1–99% confidence interval). **(B)** Horizontal projection of the location in LP of neurons from each cluster. Inset: stacked bar plot showing the fraction of cells in each cluster across LP subregions (numbers give total cell count in each bar). **(C,D,E)** Mean size tuning (C) and cumulative distributions of receptive field area (D) and patch-background index (E) for the two clusters.

### Modulation of LP neurons by motor activity

A subset of LP neurons that project to V1 are modulated by motor activity, in particular running and eye movements (Roth et al., 2016), in addition to visual stimuli. To test whether neurons across the three LP subregions were modulated by running, we compared visual responses to checkerboard and gratings stimuli during trials when mice were stationary or running (Fig. S9A-D). We found a weak but significant facilitation of visual responses during running in all LP subregions, with pLP being the least modulated. Cells that were modulated during the checkerboard stimulus tended to be similarly modulated during drifting gratings (Fig. S9D).

We also tested whether LP neurons were sensitive to eye movements by recording neural activity and eye position in the dark. Aligning the activity of LP neurons to the initiation of horizontal saccades revealed cells that were either excited or suppressed by eye movements (Fig. S9E-N). The fraction of cells that were significantly modulated during saccades was similar across LP subregions (22–24% excited, 6–10% inhibited; n=363 pLP, 472 aLP, and 49 mLP neurons from mice in which we recorded at least 10 saccades in the dark). In SC, 2/11 optotagged cells were excited and none were inhibited during saccades.

### V1 and SC input drive visual activity in separate LP subregions

Our results show that the visual response properties of pLP neurons are similar in several characteristics to those of SC neurons that project to pLP. However, it is unclear whether input from SC alone drives visual responses in pLP. In theory, the retino-geniculate-V1 pathway could provide the primary drive to pLP through both the sparse direct projection from V1 to pLP and indirect connections through higher visual areas (Fig. 2G). Direct retinal input to LP could also contribute to pLP activity, though this input is sparse and primarily from non-image forming melanopsin-expressing retinal ganglion cells (Allen et al., 2016). Conversely, though SC does not project directly to aLP, it could influence aLP through indirect pathways. To determine the relative impact of cortical and SC input to LP, we recorded spontaneous activity and visual responses of LP neurons while suppressing activity in visual cortex or SC.

To suppress activity in visual cortex, we directed blue light over V1 in VGAT-ChR2 mice (Fig. 7A; this manipulation activates inhibitory neurons which potently suppresses the local cortical network (Zhao et al., 2011)). Cortical suppression strongly reduced both spontaneous activity and responses to the checkerboard stimulus in aLP neurons (Fig. 7B-F). To a lesser extent, spontaneous activity and visual responses were also reduced in pLP neurons (Fig. 7C,D,G,H). Unlike aLP neurons, the strength of suppression of visual responses in pLP neurons depended on the stimulus condition (p = 0.019 for pLP, p = 0.83 for aLP, Kruskal-Wallis test). In pLP neurons, responses to the checkerboard patch moving over a stationary background were weakly suppressed compared to responses when the background was moving (Fig. 7H). Consequently, when cortex was inactivated, pLP neurons more strongly preferred object motion compared to background motion, becoming even more like SC neurons that project to pLP.

**Figure 7.**
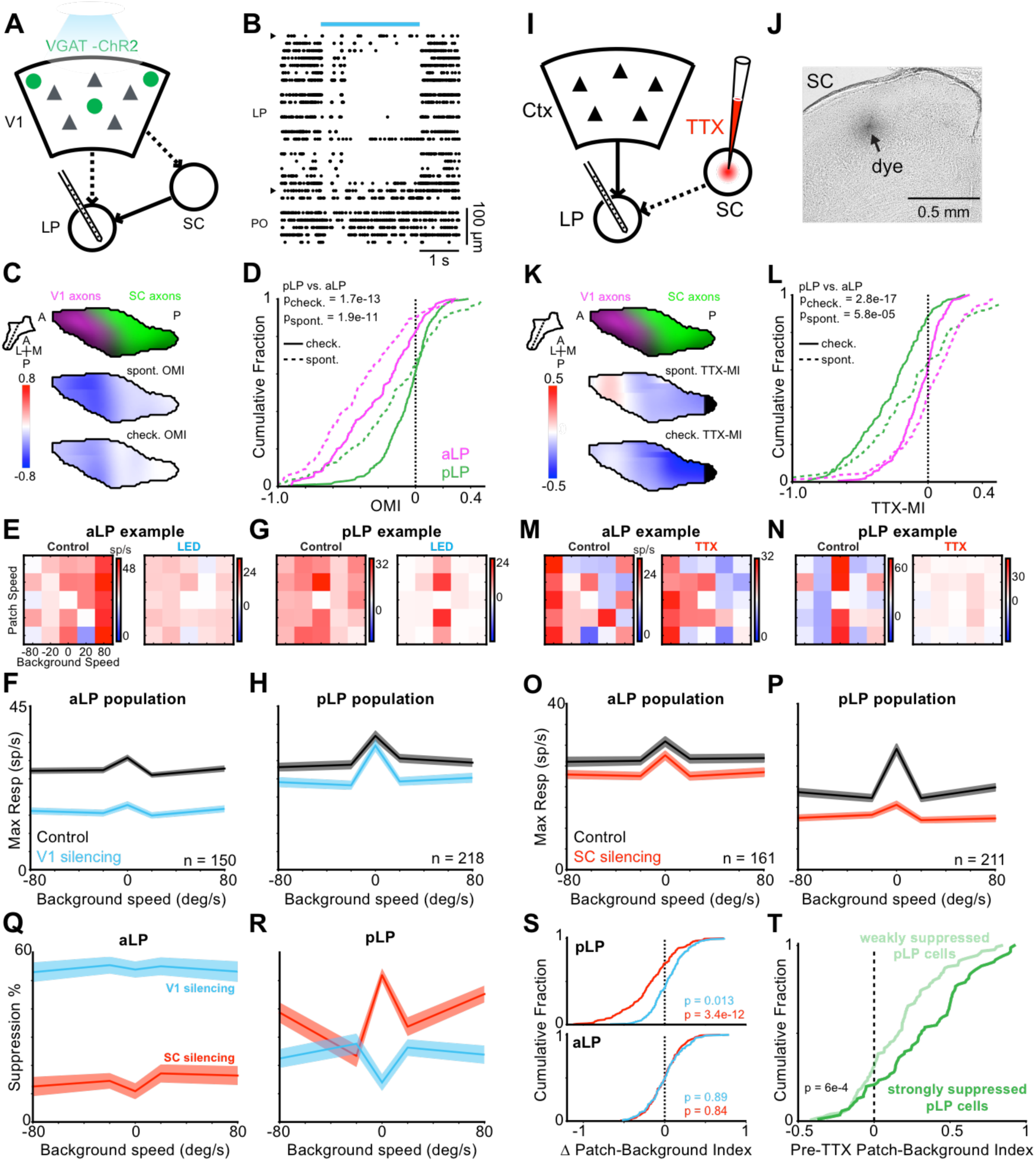
V1 and SC silencing have divergent effects on activity in anterior and posterior LP. **(A)** V1 silencing was accomplished by transcranial illumination of the cortex with blue light in VGAT-ChR2 mice. Recordings were simultaneously performed in LP. **(B)** Raster plot showing spontaneous activity of all thalamic units for one silencing trial. Blue bar indicates light delivery. Units are ordered by dorsal/ventral position. Carets demarcate LP boundaries. Units ventral of LP are in the posterior thalamic nucleus (PO). **(C)** Top: LP slice showing SC (green) and V1 (magenta) input to LP. Plane of slice indicated by dotted line in inset. Middle: Optogenetic modulation index (OMI) for spontaneous activity averaged across all units in LP. OMI is defined as (optogenetic firing rate - control firing rate)/(optogenetic firing rate + control firing rate). Bottom: OMI for checkerboard response. **(D)** Cumulative distribution of OMI for neurons in pLP (green, n=218 cells) and aLP (magenta, n=150 cells) during spontaneous activity (dotted lines) and the checkerboard stimulus (solid lines). P-values compare pLP to aLP; Wilcoxon rank-sum test. **(E)** Example patch-checkerboard matrix for an aLP neuron during control (left) and V1 silencing (right) trials. **(F)** Mean background speed tuning during control (black) and V1 silencing (blue) for aLP population. Values are maximum projections along the columns of the checkerboard response matrix averaged across cells. Shaded regions denote the SEM. **(G)**, **(H)** same as (E), (F) but for pLP. **(I)** SC silencing was accomplished by injecting TTX into the SC while recording in LP (n=211 pLP cells, 161 aLP cells). **(J)** Bright-field image confirming the deposition of dye in sSC after a TTX injection. **(K-P)** As in (C-H) for SC silencing. **(Q)** Suppression as a function of checkerboard background speed for aLP population during cortical (blue) and SC (red) silencing. **(R)** As in (Q) for pLP. **(S)** Change in the patch-background index during cortical and SC silencing for pLP (top) and aLP (bottom). A negative shift indicates a reduction in patch preference. Wilcoxon signed-rank test for shift from zero. **(T)** Distribution of patch-background index values (before TTX injection in SC) of pLP neurons that were strongly (TTX-MI<-0.33, n=86 cells) or weakly (TTX-MI>-0.33, n=125 cells) inhibited by SC inactivation. Wilcoxon rank-sum test.

The effect of cortical suppression on activity of pLP neurons could partially reflect reduced cortical input to SC neurons that project to pLP (Zhao et al., 2014) (Fig. 7A). Thus, we also measured the impact of cortical suppression on SC neurons (note that use of VGAT-ChR2 mice precluded optotagging of LP-projecting SC cells). Cortical suppression moderately reduced visual responses of neurons in the optic fiber layer of SC (SCop), where the somas of LP-projecting neurons are located (Fig. S10). Although the suppression of SCop neurons was similar in magnitude to the effect on pLP neurons, it did not significantly depend on checkerboard background speed (p = 0.86, Kruskal-Wallis test). Thus, suppression of SC activity does not fully explain the effects of cortical inactivation on pLP neurons, suggesting that the activity of pLP neurons is most likely shaped by combinations of input from both SC and cortex.

To directly assay the contribution of SC to LP visual responses, we inactivated SC with TTX while recording in LP. The relative effects of SC inactivation on aLP and pLP neurons were opposite of those observed during cortical inactivation (Fig. 7K,Q,R). Spontaneous activity and visual responses were more strongly suppressed in pLP than aLP (Fig. 7K,L). Moreover, pLP neurons lost their preference for object motion: responses to the checkerboard patch moving over a stationary background were more strongly suppressed than responses when the background was moving (Fig. 7M-S; p = 3.7e-9 for pLP, p = 0.65 for aLP, Kruskal-Wallis test). Thus, the functional properties of pLP neurons depend critically on receptive field-defining input from SC.

Silencing visual cortex or SC did not impact the running modulation of visual responses in aLP or pLP (SC silencing—ΔRMI pLP: −0.05±0.3, p=0.16; aLP: −0.02±0.2, p=0.16; V1 silencing—ΔRMI pLP: −0.006±0.2, p=0.95; aLP: −0.002±0.3, p=0.36; where ΔRMI is control – perturbation, Wilcoxon sign-rank test), suggesting that the small increase in responsiveness during running we observe across LP may reflect ascending neuromodulatory input.

pLP contains subdomains receiving bilateral or unilateral SC input (Fig. S4). Since, we did not attempt to inactivate the SC contralateral to the LP recording, the effect of silencing ipsilateral SC might be stronger in the unilateral-recipient zone than the bilateral-recipient zone. However, this was not the case, as the observed effect was moderately stronger in the bilateral-recipent zone (median checkerboard TTX-MI was −0.30 for cells in the bilateral zone (n=132) and −0.19 for cells in the unilateral zone (n=111), p=0.0032).

Responses to the checkerboard and looming stimuli, as well as functional clustering, revealed diversity among pLP neurons (Fig. 4E, 5F, 6B), suggesting some pLP neurons may depend more strongly than others on SC input. Consistent with this possibility, the pLP neurons that were most suppressed by SC inactivation showed a greater preference for object motion during control trials than less suppressed pLP neurons (Fig. 7T). Thus, functional diversity in pLP is at least in part explained by strength of input from SC.

## Discussion

We combined comprehensive anatomical circuit tracing with activity mapping across the entire LP complex in mice. By registering data from multiple modalities to a common anatomical coordinate system, we made quantitative comparisons between anatomy and functional properties. Moreover, silencing the two main LP input pathways showed divergent effects across LP. Neurons in pLP are driven by SC input and respond to looming stimuli and small, moving objects. Conversely, neurons in aLP are driven by visual cortical input and respond to large stimuli and full-field motion. A third subregion, mLP, may provide a transthalamic route for information flow from frontal/associational cortex to visual cortical regions.

### Input/output connectivity defines functionally relevant thalamic subregions

The close correspondence between anatomically and functionally defined LP subregions supports an emerging view that the fundamental units of corticothalamic computation are not individual thalamic nuclei but more precise thalamic circuits linking functionally related cortical areas (Halassa and Kastner, 2017; Shipp, 2003). A potential criticism is that the three LP subregions we define are not truly distinct, but lie along a continuum of many input/output microcircuits. However, the reversal of the elevation map, and its close correspondence to the merging of axons from SC and V1, suggests that pLP and aLP are functionally distinct zones. Moreover, mLP is defined largely by a unique projection from frontal cortex, which categorically separates it from the other LP subregions. Thus, we believe our parcellation captures three broad but functionally relevant domains of LP. Nonetheless, it should be noted that both the input and output connectivity between cortex and aLP/pLP is dense, and most of the connections between these structures that could exist are present, at least to some degree. This dense network topology could allow LP to coordinate activity across distinct cortical modules, though recent work has indicated that direct cortico-cortical connectivity is itself much more dense than previously thought. Future fine-grained investigation could reveal more precise microcircuits within each subregion.

Rodent LP was previously divided into three subregions – caudomedial (LPcm), lateral (LPl), and rostromedial (LPrm) – based on cytoarchitecture and immunocytochemical markers (Nakamura et al., 2015; Takahashi, 1985; Zhou et al., 2017). These regions differ in several ways from the LP subregions we identified based on anatomical connectivity and functional properties. LPl in previous work likely bridges pLP and aLP in our designation. Moreover, whereas we include all SC-recipient LP in pLP, previous schemes separated LP subdivisions that receive bilateral or unilateral SC input (LPcm and the posterior half of LPl, respectively). In mice, the portion of LP that receives unilateral SC input is relatively small (Fig. S4), and individual LP neurons extend their dendrites across the border between LPcm and LPl (Zhou et al., 2017). For these reasons, we did assign LP units to one or the other SC-recipient subdivision. Future studies may reveal functional differences between pLP neurons that receive bilateral or unilateral SC input.

### Transthalamic pathways mirror cortico-cortical connections

Our anatomical data suggest that indirect transthalamic pathways connecting cortical areas through LP largely mirror direct corticocortical connectivity. This parallel connectivity scheme could enable the pulvinar/LP to coordinate corticocortical communication by synchronizing activity across cortical areas (Saalmann et al., 2012). How the thalamus modulates or transforms information carried in transthalamic pathways, and how these signals differ in content or impact from those of direct corticocortical pathways, is a crucial question for understanding cortical computation.

The cortical projection to LP originates in both layer 5 (L5) and layer 6 (L6) pyramidal neurons. We found that L5 and L6 neurons from a given cortical area target similar regions in LP (Fig. S1K). It has been proposed that L5 provides a strong “driver” input to LP mediating feedforward transthalamic pathways, while L6 acts as a “modulator” in feedback pathways (Sherman, 2016). Our data suggest that both L5 and L6 participate in reciprocal corticothalamic feedback loops, similar to what has been described for frontal cortex connections with thalamus (Collins et al., 2018). Future studies could reveal unique contributions of L5 and L6 input to transthalamic pathways connecting distinct cortical areas.

### Cortical dorsal/ventral streams extend through cortico-LP and cortico-SC-LP circuits

Network analysis of cortical projections suggests the mouse visual system can be divided into ventral and dorsal streams similar to those described in other species (Wang et al., 2012). Our analysis of LP input/output connectivity indicate these cortical pathways are mirrored by cortico-thalamo-cortical projection patterns, with ventral stream areas reciprocally connected with pLP and dorsal stream areas reciprocally connected with aLP. The association of ventral stream areas with SC-recipient LP in mice represents a fundamental difference between rodent LP and primate pulvinar, in which the SC-recipient inferior pulvinar is most strongly connected with dorsal stream visual areas in the MT complex (Berman and Wurtz, 2010; Lyon et al., 2010). Interestingly, unlike dorsal stream areas in mice, the MT projection to SC is biased toward the superficial layers, similar to the SC projections from the mouse ventral stream. Thus, both species feature a loop connecting LP/pulvinar, cortex and SCs, despite the fact that the cortical component of that loop resides in the ventral stream in mice and the dorsal stream in primates. This data provides further evidence that the dorsal and ventral streams in mice are analogous but not identical to the primate visual streams, possibly indicating differences in how these species use vision.

The connectivity of mLP with frontal and associational cortices resembles the connectivity of medial and dorsomedial pulvinar in primates (Kaas and Lyon, 2007). Although most mLP neurons were visually responsive, they were less likely to have organized receptive fields, and many did not respond to sparse noise (Table S2). These data suggest that mLP neurons are involved in higher-order visual processing and do not encode simple visual features. In the future, it will be interesting to test whether these cells are particularly sensitive to behavioral context, similar to neurons in dorsomedial pulvinar (Petersen et al., 1985).

### SC provides driving, receptive field-defining input to pLP

Dual retinofugal pathways through LGN and SC are an ancient, highly conserved feature of the vertebrate visual system (Butler, 2008). In primates, the “secondary” visual pathway running through SC and pulvinar was traditionally thought to play a modulatory role in visual processing, potentially relaying saccade-modulated visual signals (Berman and Wurtz, 2011; Berman et al., 2017). Previous studies disagreed as to whether SC input could drive receptive field properties in pulvinar neurons (Bender, 1983; Casanova and Molotchnikoff, 1990), perhaps depending on species and/or where recordings were made in the pulvinar. Our data show that pLP neurons are tuned similarly to their input from SC. Moreover, silencing SC dramatically suppresses visual responses of pLP neurons and abolishes their preference for object motion (Fig. 7). Together these results demonstrate a major role for SC in driving visual responses in pLP and establish that the secondary visual pathway is an important parallel stream running alongside the geniculo-cortical pathway. Our data further suggest that this pathway is particularly important for carrying information about looming or small moving objects.

Differences between pLP and pLP-projecting SC neurons suggest that pLP is not a simple relay of SC input. In particular, receptive fields of pLP neurons are substantially larger and horizontally elongated relative to those of LP-projecting SC neurons, suggesting multiple SC neurons with horizontally displaced receptive fields might converge onto single pLP neurons. Such convergence is consistent with the topographical organization of SC axons for elevation but not azimuth in LP, and may explain the lack of an organized map of azimuth representation in pLP.

### Cortical and subcortical inputs converge in pLP

Both SC and ventral stream visual cortical areas (POR, LI) project to pLP, suggesting individual pLP neurons could integrate input from cortical and subcortical sources. Since a large majority of pLP cells were suppressed during both SC and cortical inactivation experiments, many pLP neurons likely combine input from these two sources. Interestingly, SC and cortical inactivation had differential effects on pLP excitability: cortical inactivation more strongly suppressed spontaneous than visually-evoked activity and reduced responses to background motion, while SC inactivation more strongly suppressed visually-evoked activity and abolished tuning for object motion (Fig. 7). Thus, cortical input to pLP may provide tonic modulatory drive as well as information about visual context and object identity, which could be integrated with object motion signals from SC.

The relative strength of cortical and SC input could underlie the functional diversity of pLP neurons. Indeed, we found visual tuning differences between pLP neurons that were strongly or weakly suppressed by SC inactivation (Fig. 7T). Moreover, neurons in pLP were split between two functionally defined clusters, only one of which closely resembles SC input (Fig. 6). Responses to checkerboard and looming stimuli also revealed functional diversity in pLP (Fig. 4E, 5F). The degree to which driving inputs from one or more sources (e.g. SC and/or cortical areas) converge on individual neurons is a critical outstanding question for LP and higher-order thalamic nuclei in general (Groh et al., 2014; Mease et al., 2016).

Recent studies have shown that higher order thalamus plays a fundamental role in shaping and maintaining cortical activity (Guo et al., 2017; Purushothaman et al., 2012; Schmitt et al., 2017; Zhou et al., 2016). Our comprehensive characterization of the structural and functional organization of mouse LP provides a roadmap to decipher circuit-specific mechanisms by which higher-order thalamus contributes to visual processing and behavior.

## Methods

### Mice

Rabies injections in visual cortical areas were performed at the Salk Institute and approved by the Salk Institute’s Institutional Animal Care and Use Committee (IACUC). All other experiments were performed at the Allen Institute and approved by the Allen Institute’s IACUC. Mice of either sex were used and were >70 days old on the day of electrophysiological recordings. Transgenic mice were maintained on a C57BL/6J background. The number of mice used for each experiment is indicated in Tables S1 and S2.

### Anatomy

To generate LP input maps (Fig. 1A), we utilized publicly available anterograde tracing data from the Allen Institute Mouse Brain Connectivity Database (Oh et al., 2014) (Table S1). For cortical areas, we used injections in C57BL/6J, Emx1 Cre, Rbp4-KL100 Cre, or Ntsr1-GN220 Cre mice. For SC, we used injections in C57BL/6J mice with viral expression in the superficial layers (experiments 112827164, 114754390, 126646502, 128001349, and 146078721). Experiments with injection volumes less than 0.05 mm^3^ were excluded. For injections in visual cortical areas, experiments with injection volumes greater than 0.25 mm^3^ were also excluded. For each source of input to LP, we averaged “projection energy” volumes aligned to the Allen Institute Common Coordinate Framework (CCF; 25 µm^3^ voxels) after normalizing by the brightest voxel in LP. Voxels in LP were then smoothed in three dimensions with a Gaussian kernel with standard deviation of 75 µm. Maximum-intensity projections of smoothed LP voxels in the horizontal and sagittal planes are displayed in Fig. 1A. For all analyses (e.g. comparing overlap of axons from different sources or clustering) the entire volume of voxels in LP were used.

To generate LP output maps (Fig. 2A; S2), we injected G-deleted rabies encoding fluorescent protein into each of the LP targets shown in Fig. 1 (SADΔG-EGFP or mCherry derived from the SAD B19 vaccine strain of rabies; (Wickersham et al., 2007)). For injections in visual cortical areas, injections (35–150 nL, 350 µm deep) were targeted to the center of visual areas identified by intrinsic signal imaging (Garrett et al., 2014). Anterior cingulate (ACA) injections were 0.5 mm anterior and 0.25 mm lateral from bregma and 0.5 mm deep from the pia (100–150 nL). Lateral amygdala (LA) injections were 1.3–1.8 mm posterior and 3.1–3.3 mm lateral from bregma and 3.4–4.2 mm deep from the pia (50 nL at each of three depths). Mice were perfused seven days after rabies injection. Images of 50–100 µm thick coronal sections were aligned to the CCF in three steps. (1) Images were downsampled to 25 µm pixels and the first and last sections were manually aligned to the CCF template brain. Intermediate sections were then aligned via interpolation. (2) A global affine transformation (scale, rotation, shear) was applied to each section. The transformation matrix was obtained by comparing binarized versions of the image and corresponding template section using the Open Source Computer Vision Library function findTransformECC. (3) Local warping was applied using manually defined key points on the image and corresponding template section. Delauney triangles were defined from the key points, and affine transformations were applied to warp each triangle in the image to the location of the corresponding triangle in the template section. CCF-aligned rabies-injected brains were averaged and smoothed as described for anterograde tracing data to generate the images and analysis shown in Fig. 2. Many (if not all) layer 5 visual cortical neurons that project to LP also send axon collaterals to SC. We did not observe labeled axons (or cell bodies) in SC from dG rabies injections in cortex (e.g. Fig. S2E). Thus, the signal we measure in LP from rabies injections in cortex is retrogradely labeled cell bodies and dendrites. The extent of LP that projects to V1 is probably underestimated due to the small number of injections targeted to the center of this relatively large visual area.

Input or input-output overlap was quantified by computing the normalized dot product of LP voxels in the average, smoothed volumes of anterograde (inputs) or retrograde (outputs) label (Fig. 1B, 2B). For each comparison, the dot product of the vectors of LP voxels were divided by the product of the length of these vectors.

To complement LP output volumes defined by retrograde rabies labelling, we analyzed anterograde tracing data from LP injections in the Allen Connectivity Database (Fig. S2 I-K). Injections were excluded if the injection volume was larger than 0.26 mm^3^ or smaller than 0.01 mm^3^, resulting in 13 injections. For each injection, the projection to each visual area was normalized by the total projection density to all visual areas. Output volumes were then computed for each target region by weighting the LP injection volumes by their normalized projection density to that region and smoothing with a 100 µm standard deviation Gaussian kernel in three dimensions. Due to potential viral labelling of nearby neurons in dLGN, output volumes were not calculated for V1 or lateromedial visual cortex (LM), which are known to receive strong projections from dLGN. We also used this dataset to assess the topography of LP projections to the dorsal striatum (Fig. S2N). To make this figure panel, the projection density for each LP injection was thresholded at 20 percent of the maximum voxel in the dorsal striatum (STRd) and colored according to the location of the injection centroid in the posterior-anterior axis. A subset of these LP injections were also used to quantify the laminar pattern of LP input to visual cortex (Fig. S2L,M). Five of the 13 injections were excluded due to direct labelling in dLGN. For the remaining 8 injections, the projection density to each visual area and layer were compiled into a layer x area matrix. Layer 6b data was found to include white matter axons and was excluded from analysis. To make Fig. S2M, these projection matrices were normalized by their peak value and averaged across injections. The columns of the resulting matrix (visual areas) were then ordered according to the visual hierarchy index in Harris et al. 2018 and normalized to sum to 1.

To analyze cortico-cortical connectivity (Fig. 2D), we combined injections from the Allen Connectivity Database for each cortical area. Injections were excluded if they were larger than 0.2 mm^3^ or smaller than 0.05 mm^3^ or resulted in a total density of less than 0.005 mm^3^ across all cortical targets. Injections into transgenic lines labelling inhibitory cells or with sparse cortical expression were also excluded. Each remaining injection was normalized by its total projection density across the targets listed in Fig. 2D. The strength of the projection from area X to area Y was defined as the median normalized projection density in area Y across all injections into area X. The resulting connectivity matrix was then row normalized to make Fig. 2D.

To analyze cortical projections to SC, we used the same injections used to quantify LP input but excluding injections in Ntsr1-GN220 Cre mice (which labels layer 6 cortical neurons that do not project to SC). We computed the difference of the median projection density in SCs and SCd divided by their sum. This quantity (y-axis in Fig. 2E,F) was compared to the overlap between SCs input to LP and either the LP input or output volume for each cortical area (x-axis in Fig. 2E,F, respectively).

To analyze the topography of SC and V1 input to LP (Fig. 3, S3), we primarily used anterograde tracing data from the Allen Connectivity Database. For V1, we used injections in C56BL/6J or Emx1 Cre mice with injection volumes between 0.05 and 0.25 mm^3^. To supplement the SC injection dataset (Table S1), we injected AAV-2.1-CAG-Flex-GFP and AAV-2.1-CAG-Flex-tdTomato into the SC of Ntsr1-GN209 Cre mice (Gerfen et al., 2013). In these mice, Cre expression in superficial SC is restricted to cells that project to LP (Gale and Murphy, 2014, 2018). AAV was injected iontophoretically from a pipette with tip diameter of 20 µm using 3 nA current applied 7 seconds on, 7 seconds off for 3 minutes. Both viruses were injected into each mouse, offset along the medial-lateral or anterior-posterior axis (Fig. S4J). One of the injections failed in 4 of 8 mice, resulting in 12 injections suitable for analysis. Images from these mice were aligned to the CCF as described for LP output mapping.

For each SC (n=17) or V1 (n=33) injection, we calculated injection and projection centroids in CCF coordinates using injection or projection density data (for injections from the Allen Connectivity Database) or raw fluorescence normalized to the brightest pixel in LP (for our SC injections). Injection centroids were defined as the average CCF coordinate of voxels in SC or V1 with density or normalized intensity greater than 0.25, weighted by their density/intensity. Projection centroids were similarly calculated using voxels in LP. V1 injection centroids were assigned elevation and azimuth values based on ISI maps of elevation and azimuth aligned to the CCF and averaged across mice (Fig. 3, S3). SC injection centroids were assigned elevation and azimuth values based on their position along the medial-lateral or anterior-posterior axis, respectively, which approximates the measured receptive field maps of SC neurons (Dräger and Hubel, 1976) and the topography of V1 projection centroids in SC (Fig. S3 D,K). Projection centroids were assigned the same elevation/azimuth as their corresponding injection centroids. To generate smoothed elevation and azimuth maps, projection centroid elevation or azimuth values were smoothed in three dimensions with a Gaussian kernel with standard deviation of 100 µm (SC and V1 data were smoothed separately and then combined in a single volume).

To compare LP input and output volumes to the retinotopic bias of each cortical visual area (Fig. S5), we utilized a mean elevation map for visual cortex based on ISI imaging from 14 mice (dataset from Garrett et al. 2014). For each visual area, we assigned a “measured” elevation value corresponding to the mean of the ISI elevation map over that area. We then computed two “predicted” elevations for each area by weighting the LP elevation map by either its anterograde input volume or rabies-based output volume (from Fig. 1 and 2).

### Electrophysiological recordings

Electrophysiological recordings from SC, LP, dLGN, or V1 neurons were made with Phase 2 Neuropixels probes (Jun et al., 2017) (128 channels arranged in two columns, with 20 µm between each recording site). Data were acquired at 30 kHz using the Open Ephys acquisition board and GUI (Siegle et al., 2017) and high-pass filtered (300 Hz). Mice were habituated to the recording rig for at least two weeks. On the rig, mice were head-fixed and allowed to run on a styrofoam cyclinder covered with rubber matting. On the day of recording, mice were anesthetized with isoflurane and a small craniotomy was made above the target brain region in the left hemisphere and covered with Qwikcast (World Precision Instruments). Mice recovered for at least two hours before recordings. For most mice, this procedure was repeated the following day for a new recording location (Table S2).

### Optotagging SC neurons

The SC cell type that projects to LP was identified (“optotagged”) by channelrhodopsin-2 (ChR2) activation in Ntsr1-GN209 Cre x Ai32 mice (Gale and Murphy, 2016; Madisen et al., 2012). During these recordings, a 50 µm core diameter optical fiber was inserted in the brain near the Neuropixels probe to a depth just above the SC (∼1 mm). Blue light was used to activate ChR2-expressing cells (2 s pulses, 0.5–10 mW measured from the fiber tip). Cells were considered optotagged if their mean firing rate during the last 1 s of the light pulses was greater than 5 standard deviations above the mean spontaneous rate. Spontaneous rate was calculated from the 1 s bin preceding each light pulse.

### Visual Stimuli

The mouse’s head was fixed at the center of a 24-inch diameter spherical dome (Fig. 3G). Visual stimuli were projected on the inner dome surface from two laser projectors (one on each side of the mouse) pointed at spherical mirrors placed below the running wheel. Four different visual stimuli were presented. (1) Sparse noise consisted of dark (0.6 cd/m^2^) and light (5.8 cd/m^2^) squares (5, 10, or 20° across) presented one at a time for 100 ms on a gray background (3.2 cd/m^2^). The stimulus center position for each trial was chosen pseudo-randomly from a grid of 10° spacing and ranging from −20 to 120° in azimuth (negative is left of straight in front of the mouse) and −30 to 90° in elevation (negative is below the eye). Full-field flashes were also presented. All of the stimulus positions, sizes, and contrasts were sampled once per loop in random order before initiating a new loop. (2) Two sets of moving gratings stimuli were presented. For both, the gratings filled the entire right side of the dome, drifted for 2 s, and were followed by a 1 s gray screen period before the next trial. The first set of gratings included two orientations (vertical gratings moving in the nasal-to-temporal direction and horizontal gratings moving downward), six spatial frequencies (0.01, 0.02, 0.04, 0.08, 0.16, and 0.32 cycles/°) and five temporal frequencies (0.5, 1, 2, 4, and 8 cycles/s), for a total of 60 trial types. Each trial type was presented once in random order before beginning a new loop. The second set of gratings included eight directions of motion, two spatial frequencies (0.02 and 0.16 cycles/°), and two temporal frequencies (1 and 4 cycles/s). (3) A checkerboard stimulus consisted of random patterns of dark and light 1° squares that filled both sides of the dome. The background checkerboard pattern moved such that the right and left halves converged (temporal-to-nasal motion) or diverged (nasal-to-temporal motion) at the point directly in front of the mouse. Simultaneously, a 10° patch, also consisting of a random pattern of dark and light 1° squares, moved horizontally with direction and speed independent of that of the background checkerboard pattern. Patch and background velocities were −90, −30, −10, 0, 10, 30, and 90 °/s (positive velocities are the nasal-to-temporal direction). The patch is invisible when the patch and background velocities are the same. The patch was presented at 0 and 40° elevation for LP recordings or targeted to the multi-unit receptive field elevation for SC recordings. (4) Looming stimuli were light or dark spots that expanded to simulate an object approaching at constant speed (Gabbiani et al., 1999). The expansion rate is a nonlinear function of time-to-collision and the object’s size/speed ratio, which was 10, 20, 40, or 80 ms. The initial spot radius was 0.5° and the final spot radius was 80°, which was held for 0.5 s before a 2 s gray screen inter-trial interval.

### Cortical inactivation

Cortical inactivation experiments were conducted in VGAT-ChR2 transgenic mice (Zhao et al., 2011). Before each experiment, two optical fibers (200 µm core diameter) were positioned over the skull over left V1 by stereotaxic coordinates (relative to lambda, the two fibers were 0 and 1 mm anterior and 2.8 and 2.6 mm lateral). Each fiber was coupled to a blue laser or LED calibrated to produce 2.5 mW measured at the fiber tip. During the sparse noise stimulus, control and cortical silencing conditions were interleaved in 25 trial (2.5 s) blocks. For all other stimulus protocols, control and cortical silencing trials were interleaved. Light delivery began one second before each silencing trial and ended 100 ms after the trial. Power was linearly ramped (100 ms) on and off. To allow cortex to recover after cortical silencing, four seconds were added between silencing and control trials/blocks. For all perturbation experiments, checkerboard stimulus conditions were reduced (patch and background velocities: −80, −20, 0, 20, 80 °/s) to increase trial repetitions. In separate mice not used for SC or LP recordings, we measured the lateral spread of cortical silencing using one fiber at 2.5 mW. Consistent with other studies (Guo et al., 2014), we found significant silencing at 1 mm and near complete silencing at 0.5 mm lateral from the fiber tip.

### SC inactivation

For SC inactivation experiments, we positioned a glass pipette (15–20 µm tip diameter) filled with 25 µM tetrodotoxin (TTX) into the left SC (0.5–0.7 lateral and 0–0.2 anterior from lambda and 1.2–1.5 deep from brain surface). During pipette insertion, black-white alternating flashes were played in the right hemifield and the visually-evoked potential (VEP) was monitored on an oscilloscope. The VEP became noticeably larger and more consistent at ∼1 mm depth, consistent with entry into the SC. The control dataset was collected after pipette insertion but before TTX injection. TTX was then injected by a picospritzer (2–10 PSI). During injection, the TTX pipette meniscus was video monitored to verify an injection volume of ∼50 nL. We allowed 5 minutes after injection for TTX to diffuse before collecting the TTX dataset. The checkerboard stimulus was ongoing throughout the control, injection, diffusion, and TTX epochs. At the end of most experiments we moved the Neuropixels probe from LP to SC to verify the SC silencing. No spontaneous or visual-driven spikes were observed at locations 500 µm from the injection pipette tip. Activity appeared normal 1 mm from the injection pipette tip. Evan’s blue (0.005%) was included in the pipette solution, and injection location was verified by post-hoc histology.

### Eye Tracking

Images of the right eye were acquired at 60 Hz in the dark with an infrared (IR) camera. IR LEDs were placed around the camera lens. The pupil is large in the dark and its edges are partially occluded. Rather than tracking the pupil center, we determined the horizontal position of the lateral edge of the pupil relative to a corneal reflection. Saccade times were detected automatically using a velocity threshold and manually verified.

### Data Analysis

Spike sorting was done in a two-step process. First, spikes were automatically detected and clustered using Kilosort (Pachitariu et al., 2016). Kilosort output clusters were then manually curated in phy(Rossant et al., 2016). All units passing this manual step were included for further analysis.

To register units to the CCF, brains were fixed overnight after the last experimental day and sliced on a vibratome (100 µm coronal sections). The first (posterior) and last (anterior) slices containing LP were aligned to corresponding CCF sections, and the remaining slices were aligned by linear interpolation. The probe track was then registered to the CCF by manually annotating the probe tip and the point at which it entered LP (based on DiI labeling). Individual units were then assigned positions along this track according to the distance from the tip to the channel with the maximum waveform amplitude. A similar method was used for SC recordings. Units were considered to be in LP if they fell within 100 µm of the LP border after CCF alignment (with the exception of dLGN units, which were readily identified by the sharp boundary between dLGN and LP in histological slices).

For analysis of responses to sparse noise, checkerboard, and looming stimuli, a mean spike density function (SDF) was computed for each stimulus condition by convolving the raw spike train with a Gaussian kernel (10 ms standard deviation for sparse noise or 100 ms for checkerboard and loom) and averaging across trials. The response to a given stimulus condition was taken as the peak of the mean SDF over the stimulus window (50–150 ms after stimulus onset for sparse noise, 250 ms to end of trial for checkerboard, or trial onset to collision time for loom). To quantify a cell’s responsiveness to a given stimulus, its peak response was compared to spontaneous activity. A cell was included in the analysis for a given protocol if its peak response across all conditions was greater than 5 standard deviations above the mean spontaneous firing rate. For sparse noise, the spontaneous firing rate distribution was estimated by randomly selecting n trials (where n was the number of repeats of the full stimulus set) and calculating the peak of the mean SDF from stimulus onset to 50 ms (a window that excludes the visual response). This process was repeated 200 times and the resulting 200 mean SDF peaks were taken as the spontaneous firing rate distribution. This same process was used for loom (with 100 repetitions), but the subsampled trials were restricted to those with a size to speed ratio of 80 ms (the longest trials), and the analyzed window ranged from stimulus onset to 1 second after onset. For checkerboard, the spontaneous rate was computed from trials in which the patch and background speed were 0°/s (a static random checkerboard stimulus; the middle square of the checkerboard response matrix).

For gratings stimuli, the response to a given trial condition was defined as the mean number of spikes elicited during that condition in the window from 250 ms after stimulus onset to stimulus offset. The spontaneous rate was taken from randomly interspersed trials for which the gratings stimulus was omitted (isoluminant gray screen).

Responses to sparse noise as a function of spatial position were smoothed with a two-dimensional Gaussian filter (standard deviation 10°) and then fit to a two-dimensional Gaussian to define receptive field location and area. Cells that did not respond to sparse noise (as defined above) or for which the fitting algorithm failed were not used for receptive field analysis. Receptive area is pi times the product of the major and minor radii of the fit at one standard deviation. To generate the measured LP elevation map, CCF voxels were assigned an elevation based on the mean elevation for cells assigned to that voxel (for the vast majority of voxels this was one or zero units). The resulting map was then smoothed with a Gaussian kernel (standard deviation 100 µm), linearly interpolating values for voxels without data. The same procedure was used for azimuth.

We used two measures to quantify the topographic organization of projection centroids or receptive field locations in LP. The first was the r^2^ value of a three dimensional linear fit of the elevation or azimuth assigned to the CCF coordinate of each projection centroid or receptive field. The second was Moran’s I, a measure of spatial autocorrelation, for these same values (Moran, 1950). Randomly dispersed elevations or azimuths result in Moran’s I near zero, perfect dispersion (i.e. a “checkerboard” of high and low values) results in a Moran’s I near −1, and segregated azimuths or elevations (i.e. high values in one portion of LP and low values in another) results in a positive Moran’s I. Standard error of the linear fit r^2^ and Moran’s I were calculated by randomly sampling (10000 repetitions with replacement) the data and taking the standard deviation of the 10000 resulting values.

To quantify the modulation of visual responses by motor activity in LP, we classified trials as running (mean speed > 5 cm/s) or stationary (mean speed < 1 cm/s). Because animals often spent the majority of the time in one behavioral state (usually running), we matched trial number and stimulus condition by randomly subsampling trials from the behavioral state with more trials. Cells were only included in the running analysis (Fig. S9) if they were deemed to have a significant response to the visual stimulus before separating trials by motor activity. The running modulation index was defined as (run + stationary)/(run – stationary), where run and stationary refer to the mean visual response across all stimulus conditions during running and stationary epochs respectively.

To quantify the effects of silencing cortex or SC on LP visual responses, we defined optogenetic and TTX modulation indices (OMI and TTX-MI) as (perturbation– control)/(perturbation+control), where “perturbation” and “control” were the peak visual response for the checkerboard stimulus averaged over the perturbation and control trials respectively. For spontaneous activity, the central square of the checkboard response matrix was used (static background stimulus). Suppression tuning curves were calculated for the best response at each background speed (maxima along the columns of the checkerboard response matrix). Only cells with significant responses to the checkerboard stimulus during control trials were analyzed.

All statistical comparisons were non-parametric, two-sided tests. For all comparisons, the test and p-value are given in the Results, figure and legend, or Table S3.

## Supporting information

Anatomy Injection IDs

## Acknowledgements

We thank Dan Denman and Josh Siegle for advice on Neuropixels recordings and Marty Mortrud for help with iontophoretic virus injections. This work was supported by the Allen Institute for Brain Science. We thank the Allen Institute founder, Paul G. Allen, for his vision, encouragement, and support.

## Author contributions

CB and SDG conceived the project, performed experiments, analyzed data and wrote the manuscript. MEG and MLN performed rabies tracing experiments under EMC’s supervision. SRO supervised CB and SDG and helped write the manuscript. GJM helped conceive the project. All authors contributed to editing the manuscript.

## Competing interests

The authors declare no competing interests.

---

**Figure S1.**
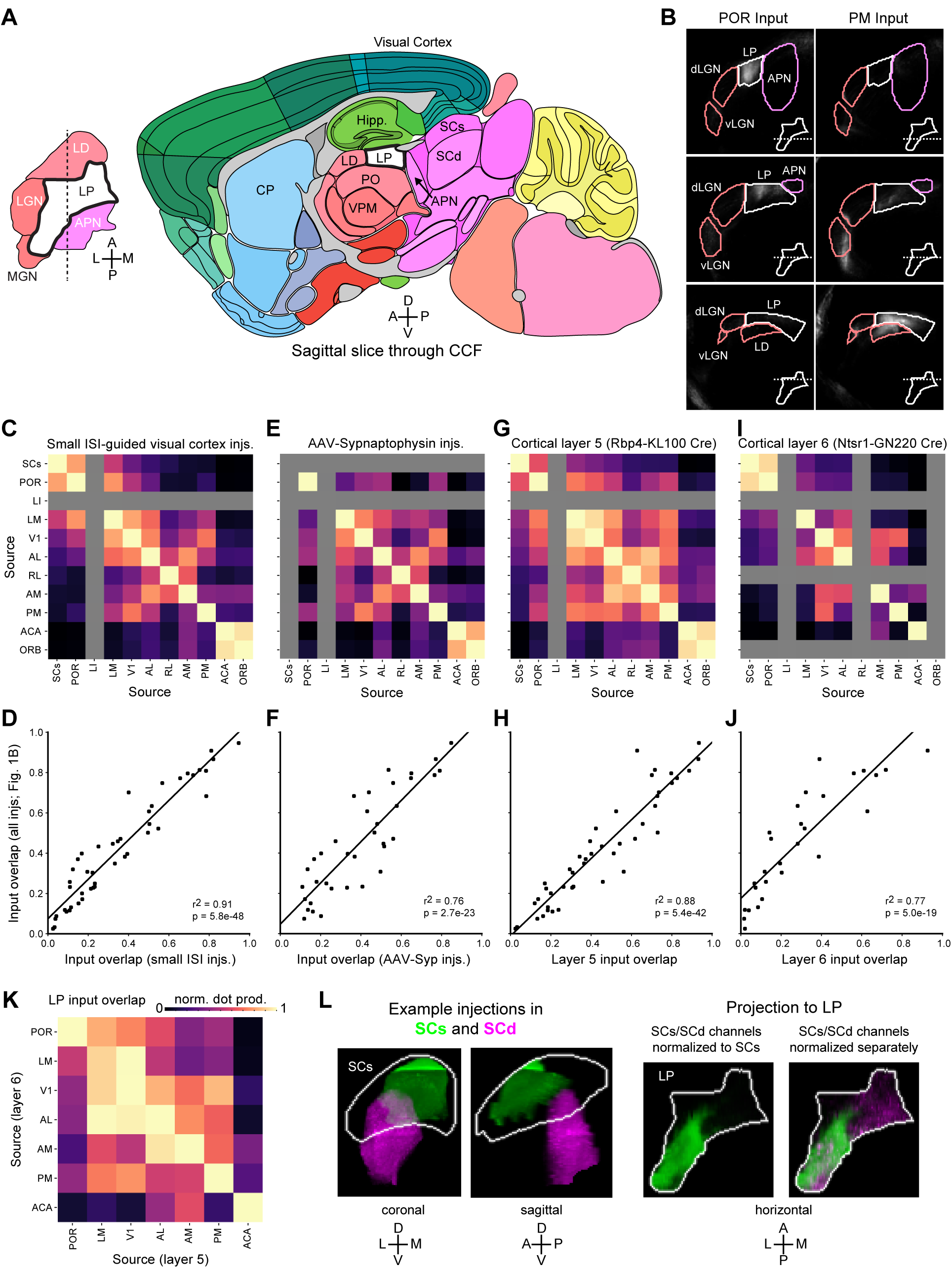
Supplementary information related to LP input. **(A)** Left: horizontal project of LP and surrounding structures using borders defined by the Allen Institute Common Coordinate Framework (CCF). Right: sagittal brain slice through LP at the location indicated by the dashed line in the horizontal projection. **(B)** Example coronal slices of the volumes used to make horizontal and sagittal projections in Fig. 1A for POR and PM input to LP. Anterior-posterior position of each slice is indicated by a dashed white line over a horizontal projection of LP borders. Axon labeling within the borders of LP was used to make the maximum-intensity projections shown in Fig. 1A. Axons also extend into other visually responsive areas that surround LP (e.g. LD). **(C)** Overlap (normalized dot product) of input to LP from different sources (similar to Fig. 1B) using a subset of experiments in which visual cortical areas were targeted by intrinsic signal imaging (ISI) and the maximum injection volume was 0.13 mm^3^ rather than 0.25 mm^3^ (Table S1). Gray boxes indicate no available data. **(D)** Scatter plot comparing the non-diagonal elements of the matrix in (C) to the same data for all injections (Fig. 1B). **(E-J)** Same as (C,D) for injections of virus encoding synapse-localized GFP (AAV-Synaptophysin-GFP) or injections in Rbp4-KL100 Cre or Nstr1-GN220 Cre mice (labeling layer 5 or 6 cortical neurons, respectively). **(K)** LP projection overlap matrix for injections in Rbp4-KL100 or Ntsr1-GN220 Cre mice. The Allen Connectivity Database does not include injections in these Cre lines for all of the cortical areas shown in Fig. 1 (Table S1). **(L)** Example AAV injections from the Allen Connectivity Database that primarily overlap superficial (SCs, green; experiment126523066) or deeper (SCd, magenta; 114754390) layers of the superior colliculus. Images of axons in LP are normalized to the brightest voxel within the volume of LP for both injections (left) or separately for each injection (right). The magenta axon labeling is not visible in the left image of LP because input from SCd is much sparser than input from SCs.

**Figure S2.**
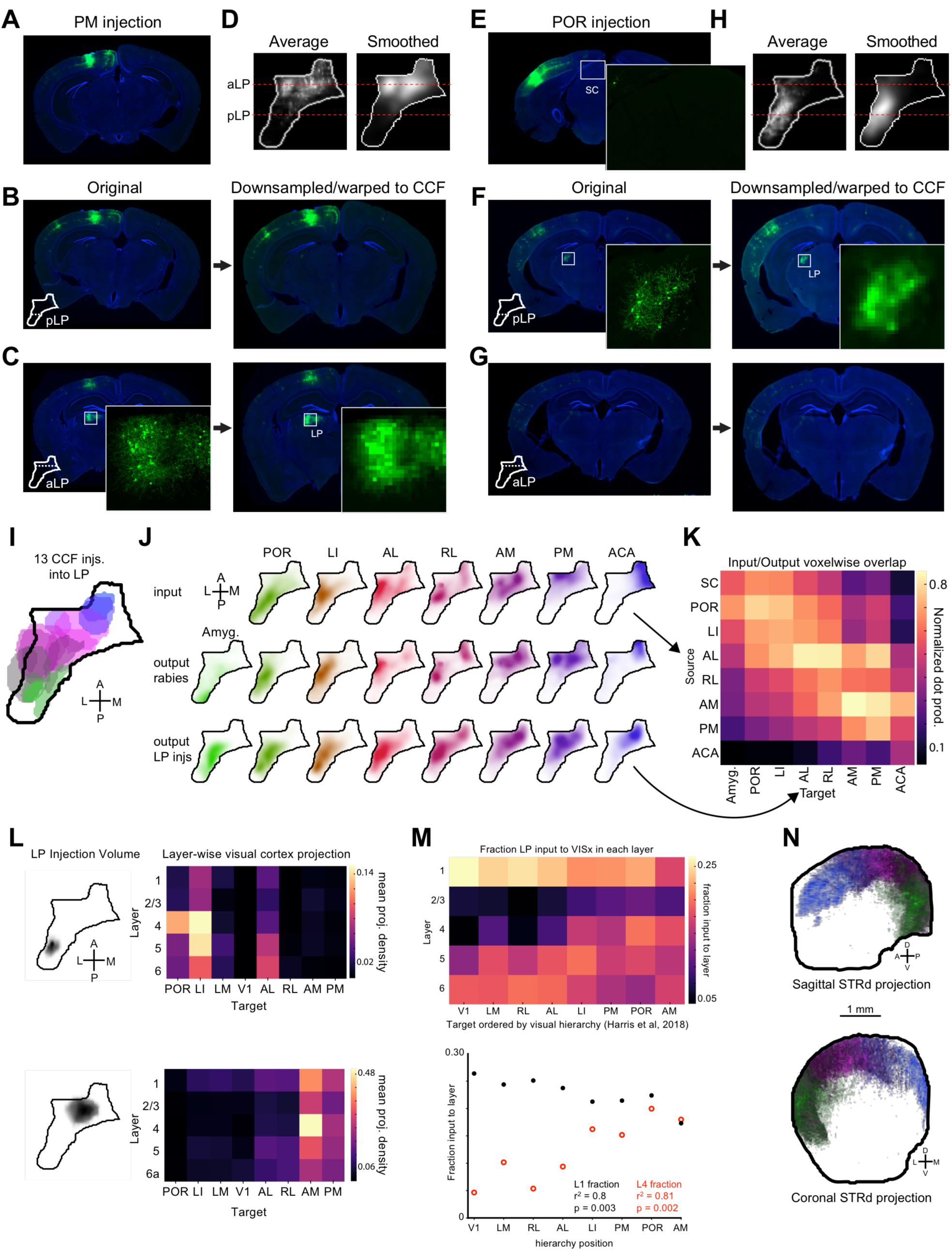
Supplementary information related to LP output. **(A)** Example G-deleted rabies-GFP injection in cortical area PM (identified by intrinsic signal imaging). **(B,C)** Retrogradely labeled cells in posterior and anterior LP from the injection shown in (A). Images were warped to the Allen Institute Common Coordinate Framework for comparisons across experiments (Methods). **(D)** Fluorescent signal in LP voxels were averaged across all PM injections and then smoothed in three dimensions. Horizontal projections of this data are shown before and after smoothing. **(E-H)** Same as (A-D) for an example injection in POR. Insets in (C) and (F) show retrogradely labeled somas and dendrites in LP. Inset in (E) shows lack of visible axon labeling in SC following rabies injections in visual cortex. **(I)** Injection volumes for 13 LP injections in the Allen Connectivity Database. **(J)** Horizontal projections of input volumes (top), rabies-based output volumes (middle, as in Fig. 1), and output volumes based on LP injections in (C). To make each injection-based output volume, the injection volumes in (C) were weighted by their projection density in the target structure of interest (Methods). V1 and LM were excluded due to possible contamination from dLGN labelling. **(K)** Input/output overlap matrix (as in Fig. 2B) comparing input volumes to LP injection based output volumes. **(L)** Laminar pattern of projections from LP to cortical areas for two example LP injections. Left: Horizontal projections of LP injection volumes. Right: Mean projection density matrices to each layer and visual cortical area. **(M)** Top: heatmap indicating the proportion of LP input to each layer for every visual area. Columns sum to one. Areas are ordered according to position in visual hierarchy. Bottom: scatterplot of data from layer 1 (black) and layer 4 (red) rows in above matrix. **(N)** Topography of LP projections to dorsal striatum shown in sagittal (top) and coronal (bottom) views for same LP injections as in (I). Projections are colored according to (I).

**Figure S3.**
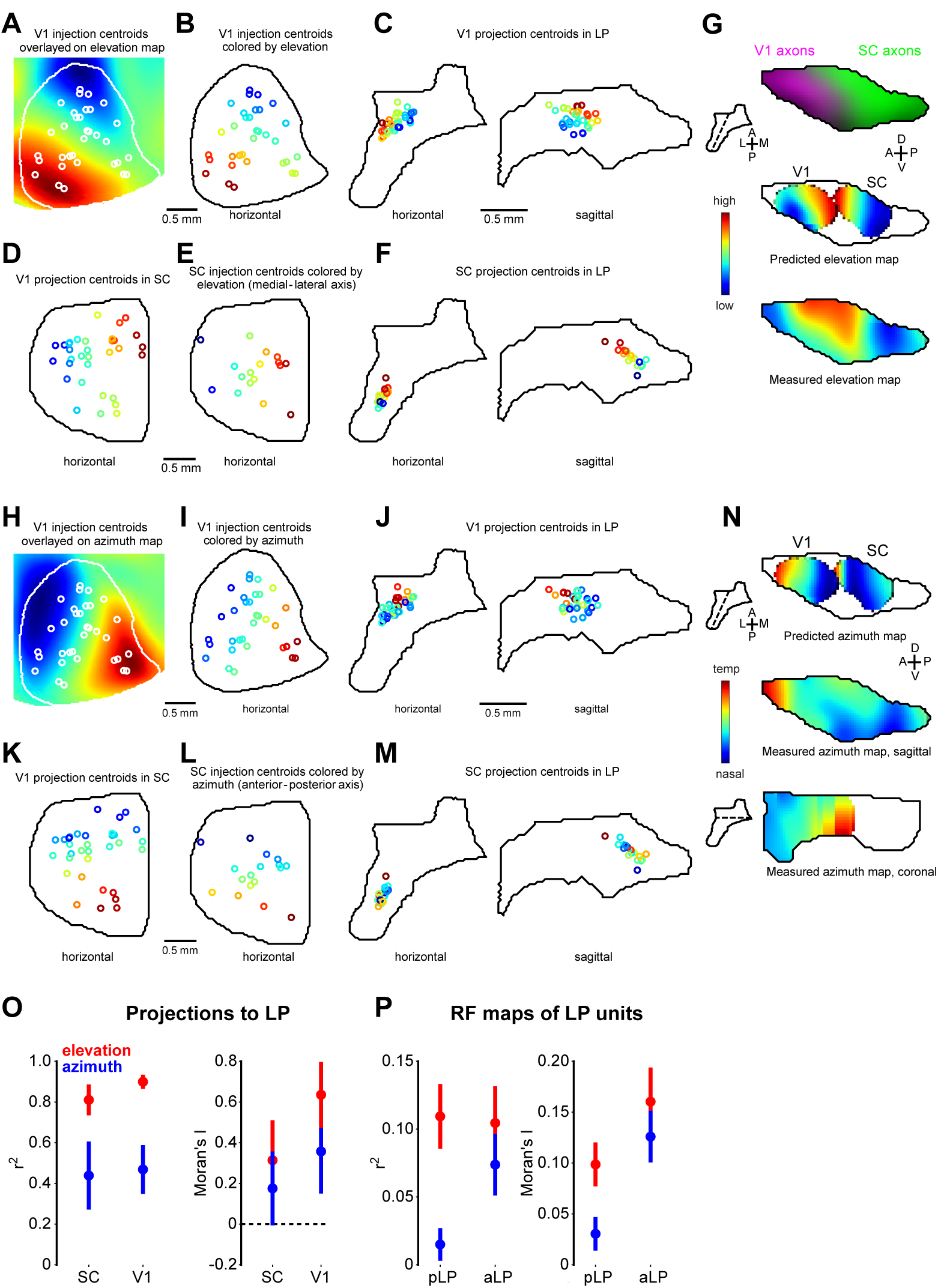
Prediction of LP elevation and azimuth maps by topography of V1 and SC input. **(A)** Mean elevation map of V1 (white outline) with location of V1 injections from the Allen Connectivity Database superimposed (white circles indicate injection centroids). **(B)** V1 injections colored by assigned elevation according to map in (A). **(C)** V1 projection centroids in LP for injections in (B) colored by assigned elevation. These centroids were smoothed to create the V1 predicted elevation map in (G). **(D)** V1 projection centroids in SC to demonstrate that the SC elevation gradient is organized along the medial-lateral axis. **(E)** Location of SC injections colored by elevation inferred from medial-lateral coordinate. **(F)** SC projection centroids in LP. **(G)** Top: LP slice showing SC (green) and V1 (magenta) input to LP. Plane of slice indicated by dotted line in inset. Middle: Predicted LP elevation map based on anatomical V1 input (left, from (B, C)) or SC input (right, from (E, F)). Bottom: Composite elevation map for all LP cells. **(H-N)** as in (A-G) but for azimuth. **(O)** Left: r^2^ values for linear fit of LP elevation (red) and azimuth (blue) projection centroids. Right: Moran’s I (see Methods) calculated for LP projection centroids. **(P)** As in (O) for measured receptive field elevations and azimuths in pLP and aLP. Error bars represent standard error estimated from random sampling.

**Figure S4.**
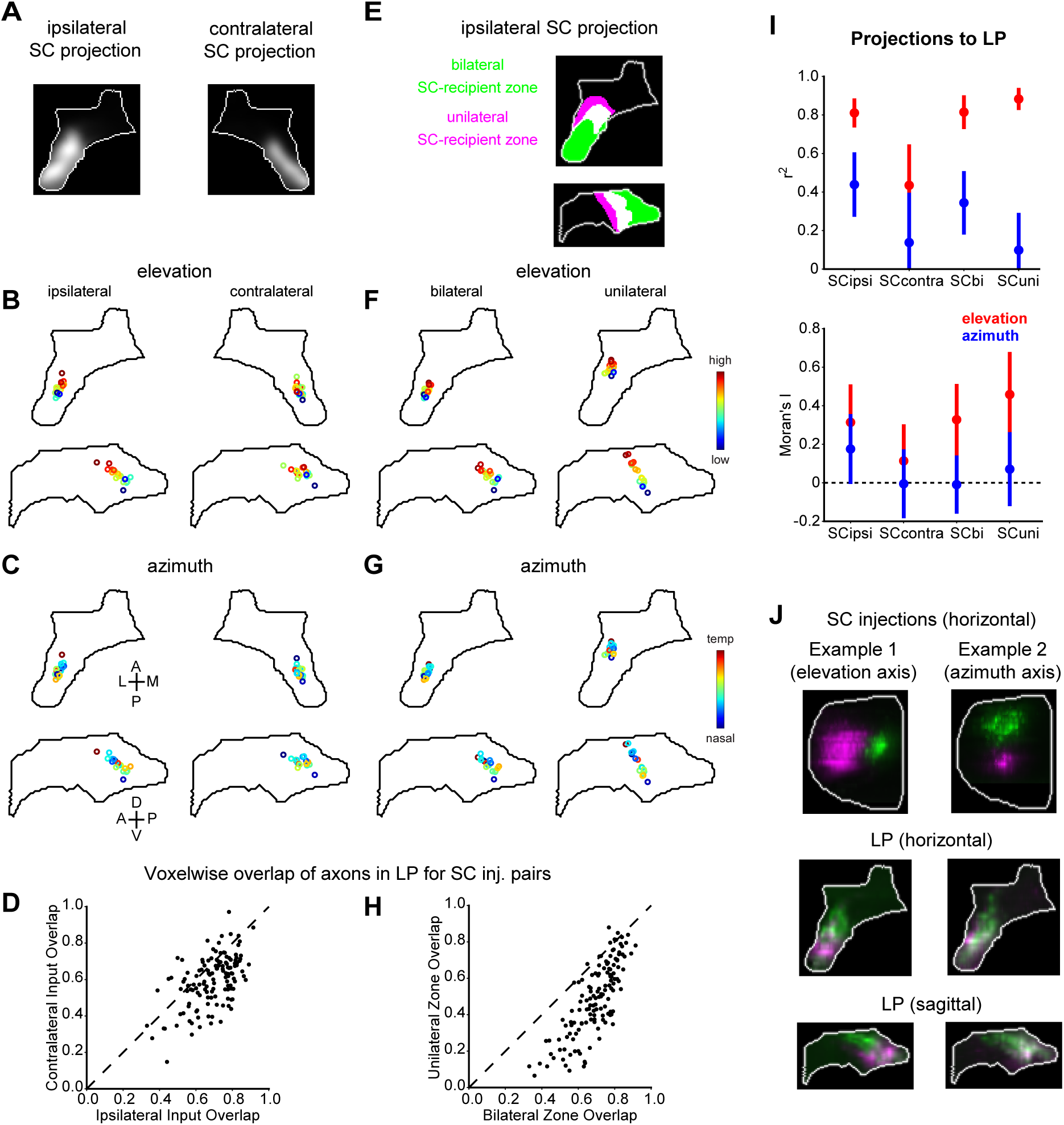
Topography of SC axons in unilateral and bilateral SC-recipient portions of LP. **(A)** Horizontal projections of average fluorescent intensity of ipsilateral and contralateral SC input to LP (as in Fig. 1A). **(B)** Ipsilateral and contralateral LP projection centroids for all SC injections colored by the relative elevation represented at the location of the injection centroid. **(C)** Same as (B) for azimuth. **(D)** Voxelwise overlap (normalized dot product) of SC axons in the bilateral SC-recipient portion of ipsilateral or contralateral LP for all pairs of SC injections. **(E)** Binarized versions of the data shown in (A) were used to define the portions of LP receiving bilateral (green) or unilateral (magenta) SC input. These subdivisions are mutually exclusive in the 3D volume but overlap (white pixels) in parts of the horizontal and sagittal projections of LP. **(F)** Projection centroids, colored by assigned elevation as in (B), of ipsilateral SC axons in the bilateral or unilateral SC-recipient portions of LP. **(G)** Same as (E) for azimuth. **(H)** Overlap of ipsilateral SC axons in the bilateral or unilateral SC-recipient portion of LP. **(I)** Top: r^2^ values for linear fit of SC elevation (red) and azimuth (blue) projection centroids in LP. Bottom: Moran’s I for SC projection centroids in LP. Data is shown for all of ipsilateral LP (SCipsi), contralateral LP (SCcontra), and the bilateral (SCbi) and unilateral (SCuni) SC-recipient portions of ipsilateral LP. **(J)** Examples of SC injections separated on the axis along which receptive field elevation (example 1; medial-lateral) or azimuth (example 2; anterior-posterior) varies. For each example, two injections were made in the same brain using virus coding for different fluorophores. The top images show the injection sites and the images below show horizontal and sagittal projections of axons in LP.

**Figure S5.**
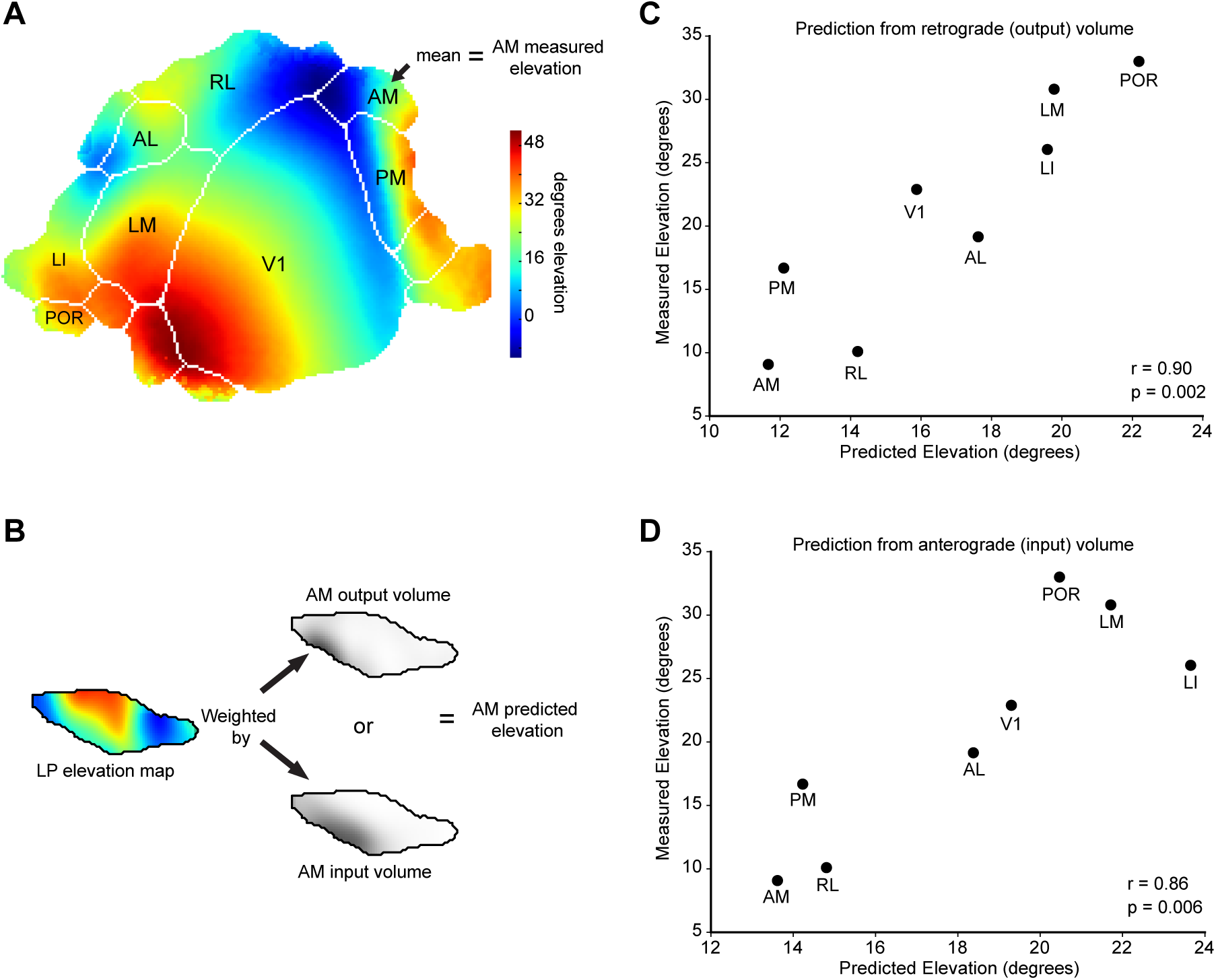
LP input/output projection volumes for eight visual cortical areas reflect biases in retinotopic coverage. **(A)** Average elevation map determined by ISI imaging across 14 animals (Garrett et al., 2014). The mean elevation across each area was assigned as the “measured” elevation in (C) and (D). **(B)** For each area (AM used here as example), a “predicted” elevation was assigned by weighting the LP elevation map by its retrograde output projection volume (used in (C)) or anterograde input projection volume (used in (D)). **(C,D)** Scatterplots comparing measured to predicted elevations based on retrograde volumes (C) or anterograde volumes (D).

**Figure S6.**
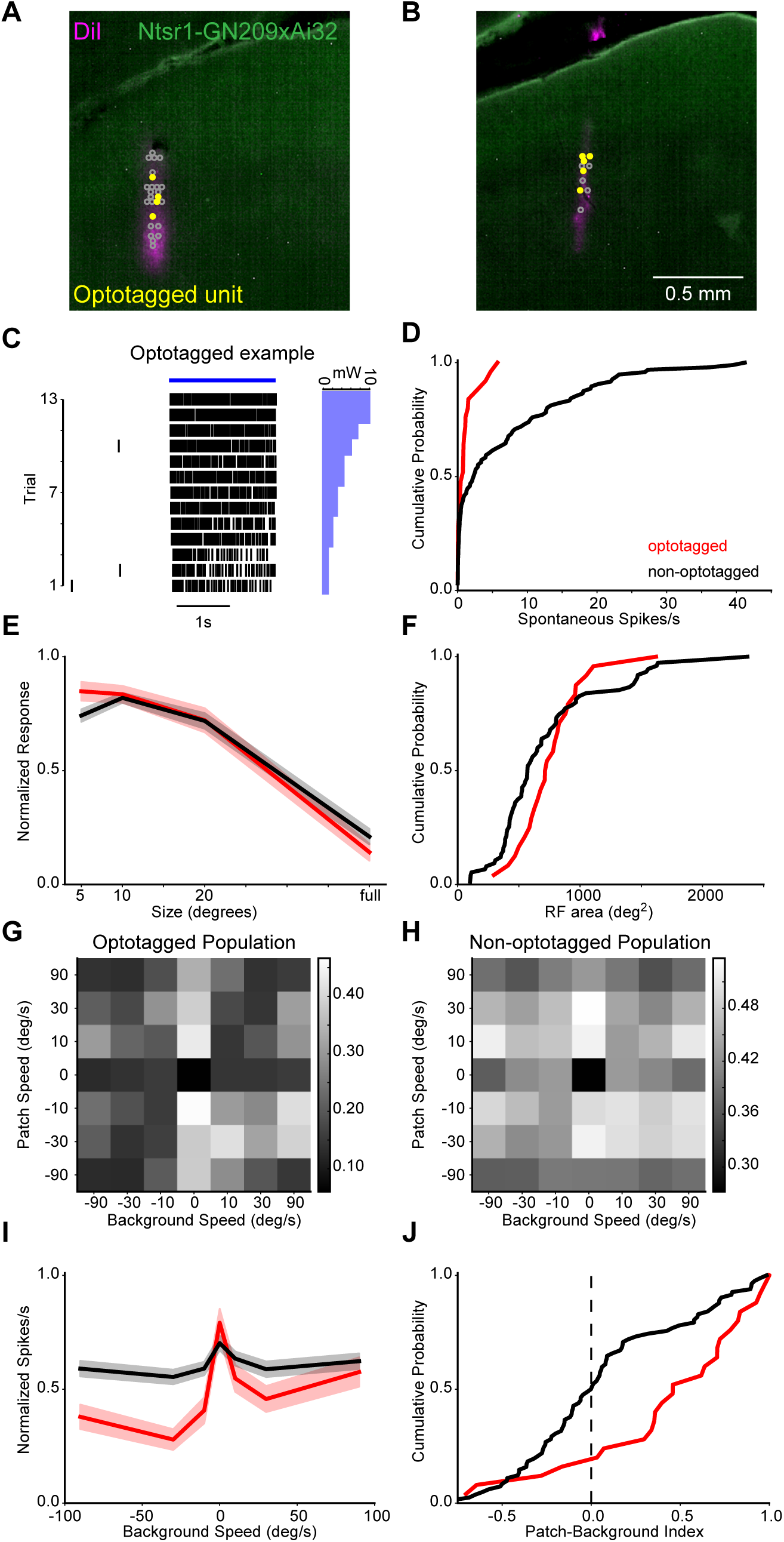
Optotagging identifies LP-projecting cells in superficial SC. **(A, B)** Coronal SC sections from two optotagging experiments showing distribution of optotagged (filled yellow circles) and non-optotagged (open gray circles) units in SC. **(C)** Raster showing response of example optotagged cell over 13 laser stimulation trials. Bar plot on right indicates laser power for each trial. **(D, E, F)** Cumulative distributions of spontaneous activity, population size tuning curves, and cumulative distributions of receptive field area for optotagged (red) and non-optotagged cells (black). **(G)** Population normalized checkerboard response matrix for optotagged SC cells. **(H)** Same as (G) for non-optotagged SC cells. **(I, J)** Population checkerboard background speed tuning curves (column maxima) and cumulative distributions for patch-background index for optotagged (red) and non-optotagged cells (black).

**Figure S7.**
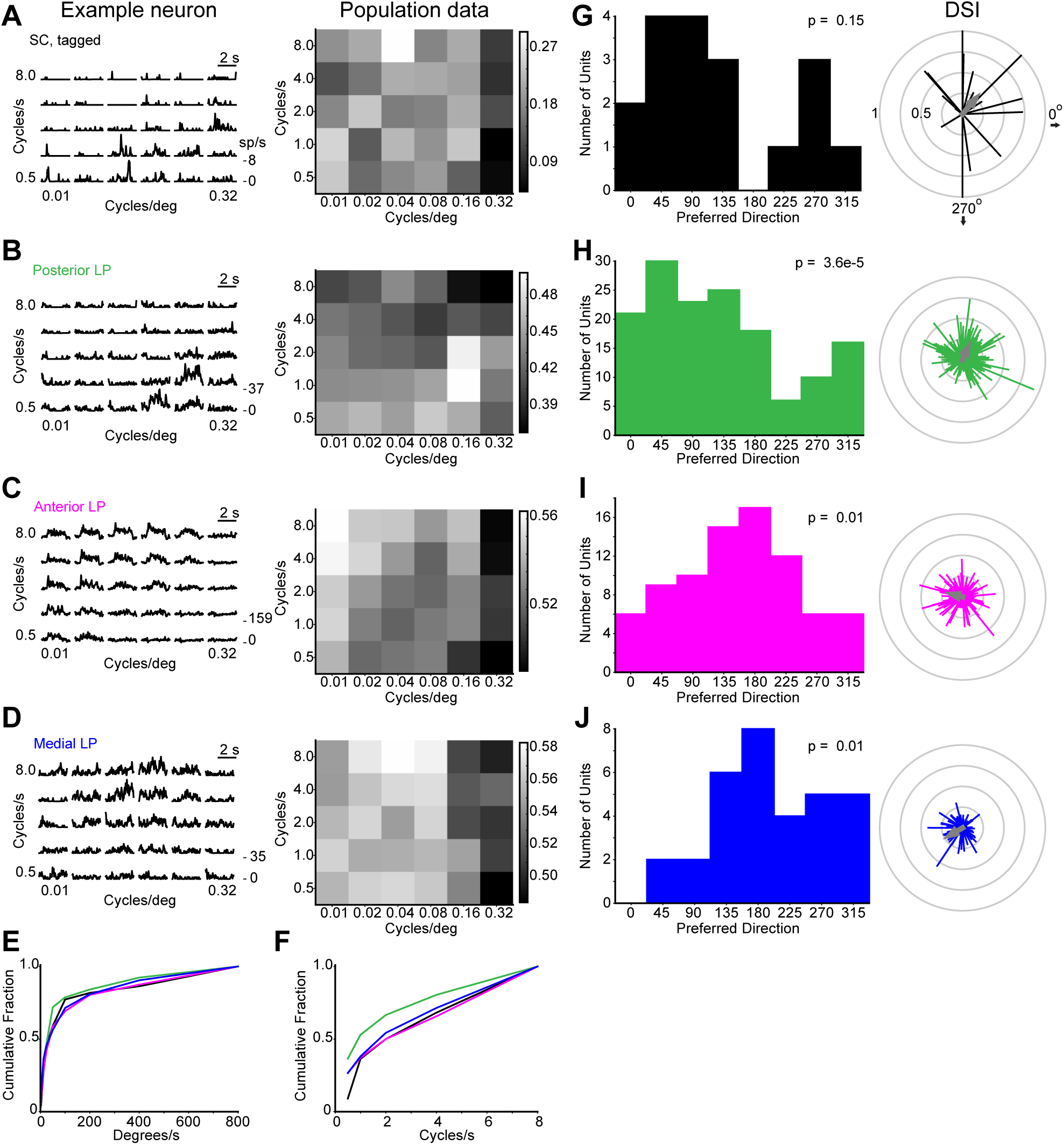
Comparison of spatio-temporal frequency and direction tuning in SC and LP subregions. **(A)** Spike density functions of an example optotagged SC neuron (left) and mean population responses (right) to drifting gratings of varying spatial and temporal frequency. **(B, C, D)** Same as (A) for pLP, aLP and mLP neurons. **(E, F)** Cumulative distributions of preferred speed and temporal frequency for SC and LP subregions. **(G)** Left: histogram of preferred directions for optotagged SC cells with direction selectivity (DSI, 1-circular variance) greater than 0.15. Right: polar plot representation of direction preference for SC cells. Each cell contributes one vector. Direction of vector indicates preferred direction and length indicates DSI. Gray arrow represents vector sum of population. **(H-J)** Same as (G) for LP subregions. P values in (G-J) calculated by Rayleigh’s test for circular uniformity.

**Figure S8.**
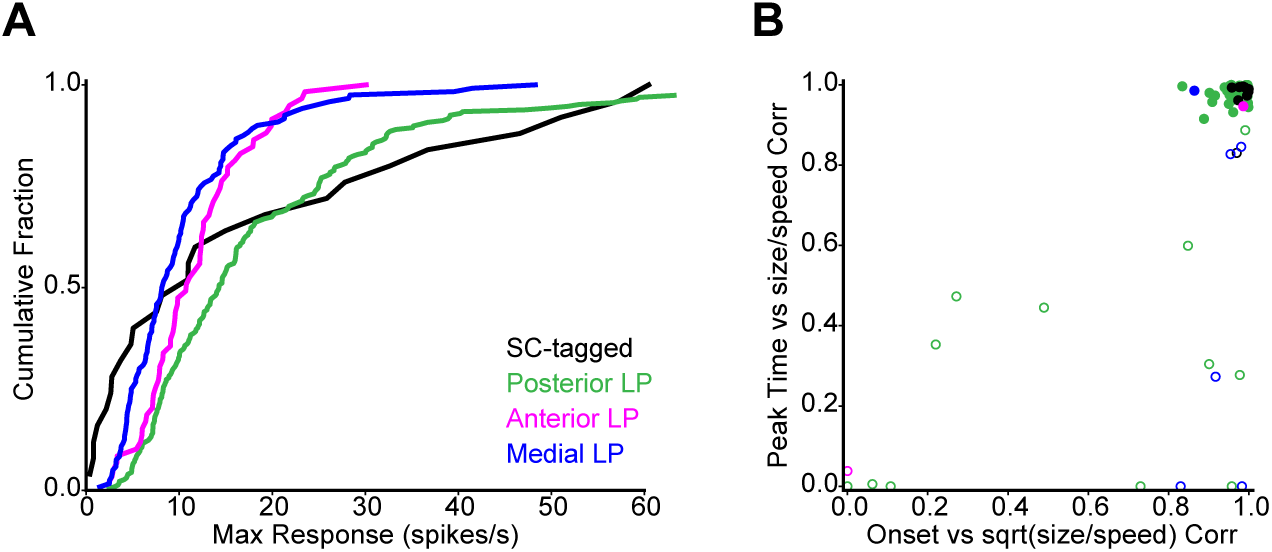
Classification of loom response types. **(A)** Cumulative distribution of maximum firing rate response to looming stimuli for cells in SC and LP subregions. **(B)** Scatterplot of correlation between peak response time and size to speed ratio (y axis) and correlation between response onset and the square root of the size to speed ratio (x axis). Cells were included only if they had a response greater than 3 standard deviations above baseline) for each size to speed ratio. Filled circles are cells that were classified as η-type. This data suggests that the majority of cells with a significant loom response display η behavior (upper right quadrant) and not two other common loom response patterns, ρ (lower right) or τ (lower left)(Sun and Frost, 1998).

**Figure S9.**
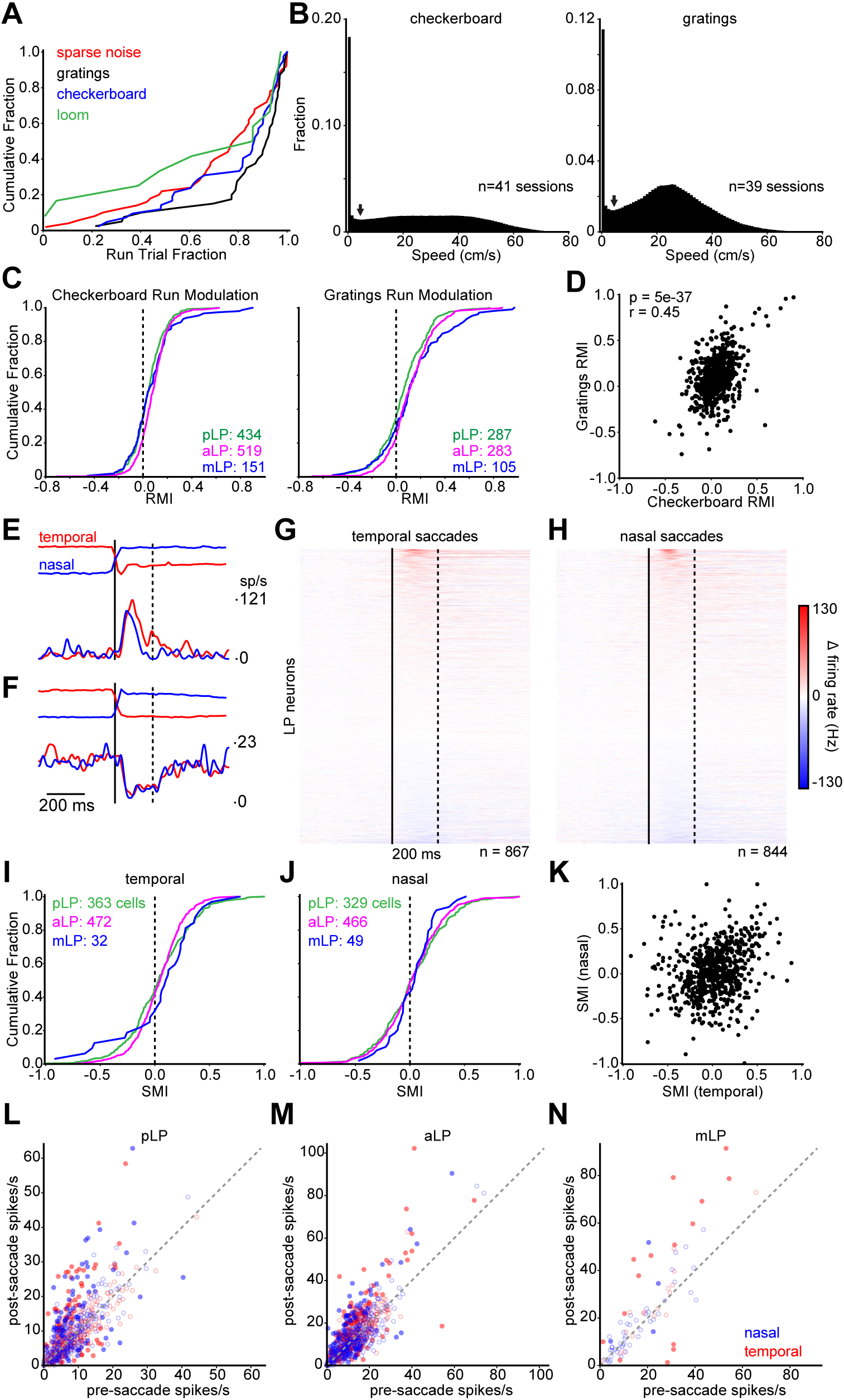
Motor activity weakly modulates cells across LP. **(A)** Cumulative distribution of run trial fraction across all recording sessions for four visual stimulus protocols. **(B)** Histograms of running speed across all sessions and mice for checkerboard and gratings protocols. The first bin defines “stationary” epochs (< 1 cm/s). Arrows indicate run threshold (5 cm/s). **(C)** Cumulative distributions of the run-modulation index (RMI)—defined as (run – stat)/(run + stat) where run and stat are the mean visual response during running and stationary trials respectively—during the checkerboard stimulus and drifting gratings. **(D)** Scatterplot comparing RMI during gratings and checkerboard for all LP cells with a significant response to both protocols (Methods). **(E)** Example unit excited by eye movements. Top traces are average eye trajectories for nasal (blue) and temporal (red) saccades. Bottom traces are mean SDFs triggered on saccade onset (solid line). The analysis period for (G-N) is bracketed by solid and dotted lines. **(F)** Same as (E) for example unit suppressed by eye movements. **(G)** Heat plot showing change in firing rate aligned to saccade onset (temporal direction) for LP cells. Only cells for which there were at least 10 saccades were included (n=867). Solid and dotted lines bracket analysis window as in (E, F). **(H)** Same as (G) for nasal saccades (n=844). **(I)** Cumulative distribution of saccade modulation index (SMI; difference between firing rates before and after the saccade divided by the sum of these rates) for cells in each LP subregion (colors as in (C)), calculated for eye movements towards the temporal visual field. **(J)** Same as (I), but for nasal saccades. **(K)** Scatterplot comparing SMI for temporal and nasal saccades. Every point represents one cell. **(L, M, N)** Scatterplots of firing rate before and after saccade onset for cells in pLP, aLP and mLP. Every cell contributes two points to a given plot corresponding to nasal (blue) or temporal (red) saccades. Filled circles denote cells with statistically significant modulation (Wilcoxon signed-rank test, p<0.01).

**Figure S10.**
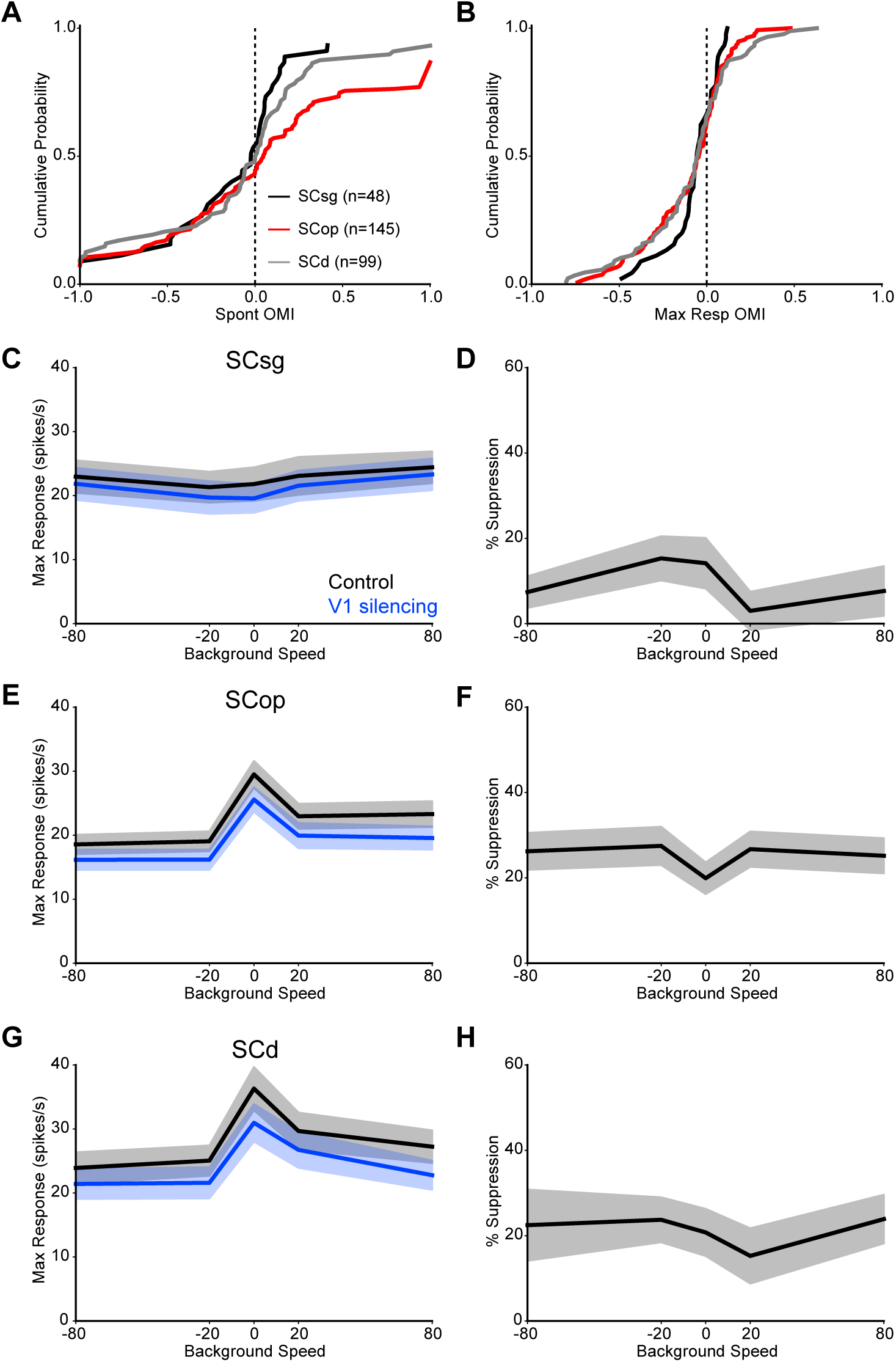
Cortical silencing moderately suppresses visual responses of SC neurons. **(A)** Cumulative distribution of the optogenetic modulation index (OMI) during spontaneous activity for SC cells in the superficial gray (SCsg), optic fiber layer (SCop), and deeper layers (SCd). **(B)** As in (A) but for the maximum response to the checkerboard stimulus. **(C)** Background speed tuning for the checkerboard stimulus (max response for each column of the checkerboard response matrix) during control and cortical silencing trials for cells in the SCsg. **(D)** Suppression during cortical silencing across checkerboard background speeds. **(E, F, G, H)** Same as (C, D) but for cells in SCop and SCd

**Supplementary Table 1:**
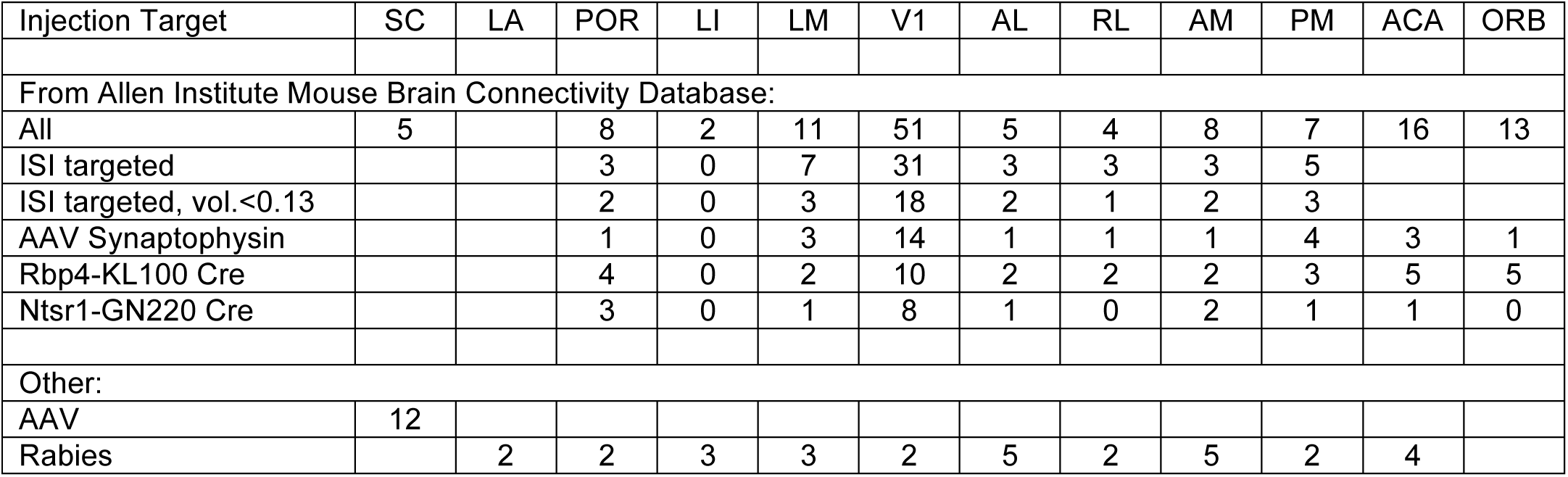
Number of injections used for anatomy experiments. In most cases, one injection was made in a mouse. For our AAV injections in SC, 8 of the 12 injections were dual injections of two different AAVs in the same hemisphere of 4 mice (see Methods). For Allen Institute Mouse Brain Connectivity Database experiments, “all” refers to experiments in wildtype, Emx1 Cre, Nstr-GN220 Cre, or Rbp4-KL100 Cre mice using AAV-GFP or AAV-Synaptophysin-GFP. For visual cortical areas, most injections were targeted using intrinsic signal imaging (ISI), and the maximum injection volume used was 0.25 mm^3^ or 0.13 mm^3^ (“small”). These data were combined for LP input mapping shown in Fig. 1. Data from specific subsets of experiments (small ISI-targeted injections, AAV-Synaptophysin-GFP, Rbp4-KL100 Cre, Nstr1-GN220 Cre) are shown in Fig. S1. For rabies injections, visual cortical areas were targeted using ISI for all experiments.

**Supplementary Table 2:**
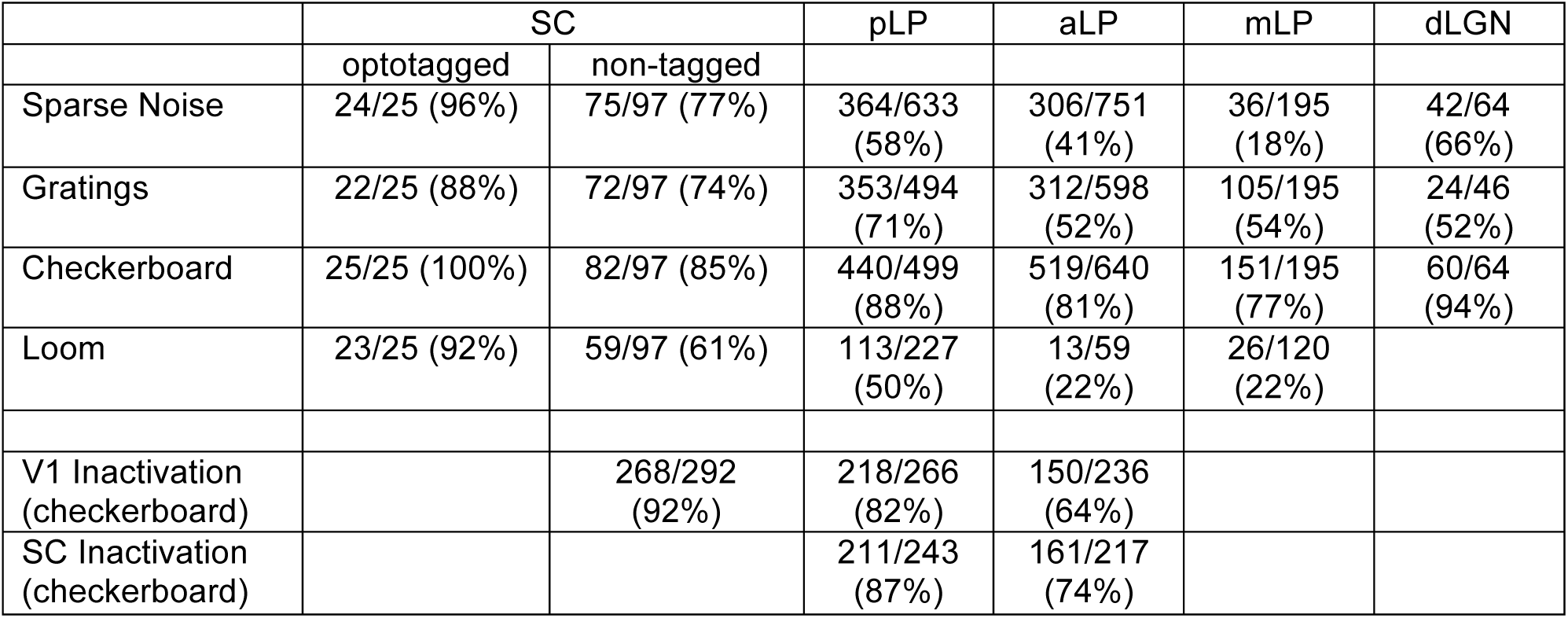
Number of cells recorded during electrophysiology experiments. Values indicate the fraction of cells that responded (see Methods) to each of the visual stimulus types shown in the left column, or to the checkerboard stimulus for V1 and SC inactivation experiments. For the sparse noise stimulus, a spatial receptive field that could be fit to a two-dimensional Gaussian was also required to include a cell in the numerator. Only a subset of the stimuli were presented during some experiments. SC optotagging experiments were in Ntsr1-GN209 Cre x Ai32 mice. Cortical inactivation experiments were in VGAT-ChR2 mice. The number of mice used for control experiments were 5 (SC), 30 (LP) and 3 (dLGN); for V1 inactivation 5 (SC) and 6 (LP); and for SC inactivation 8 (LP). In some cases we recorded twice from the same mouse on consecutive days. The number of recording sessions for control experiments were 8 (SC), 50 (LP), and 3 (dLGN); for V1 inactivation 9 (SC) and 11 (LP); and for SC inactivation 10 (LP).

|  | SC |  | pLP | aLP | mLP | dLGN |
| --- | --- | --- | --- | --- | --- | --- |
|  | optotagged | non-tagged |  |  |  |  |
| Sparse Noise | 24/25 (96%) | 75/97 (77%) | 364/633<br>(58%) | 306/751<br>(41%) | 36/195<br>(18%) | 42/64<br>(66%) |
| Gratings | 22/25 (88%) | 72/97 (74%) | 353/494<br>(71%) | 312/598<br>(52%) | 105/195<br>(54%) | 24/46<br>(52%) |
| Checkerboard | 25/25 (100%) | 82/97 (85%) | 440/499<br>(88%) | 519/640<br>(81%) | 151/195<br>(77%) | 60/64<br>(94%) |
| Loom | 23/25 (92%) | 59/97 (61%) | 113/227<br>(50%) | 13/59<br>(22%) | 26/120<br>(22%) |  |
| V1 Inactivation<br>(checkerboard) |  | 268/292<br>(92%) | 218/266<br>(82%) | 150/236<br>(64%) |  |  |
| SC Inactivation<br>(checkerboard) |  |  | 211/243<br>(87%) | 161/217<br>(74%) |  |  |

**Supplementary Table 3:**
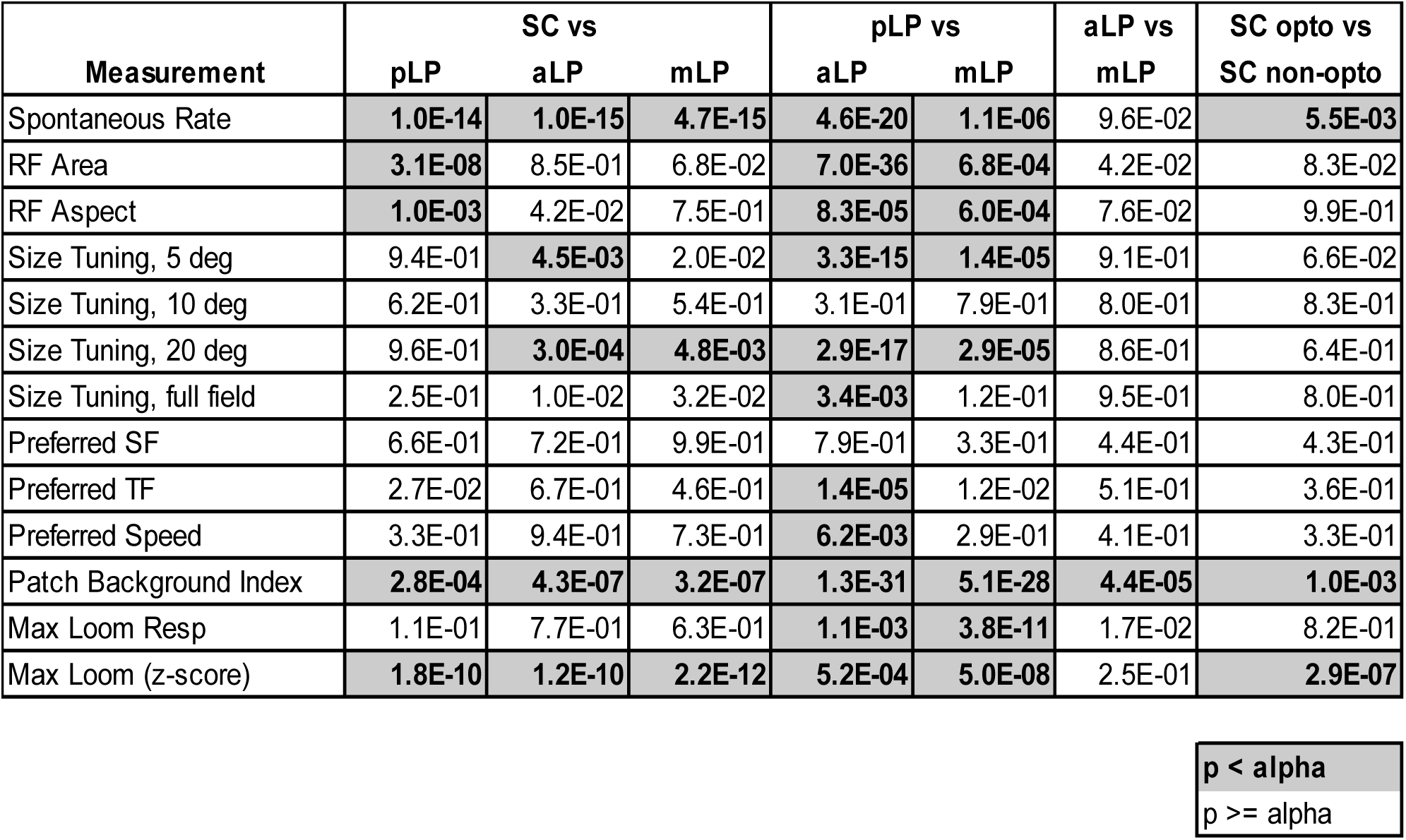
Statistical comparisons for neurons in different recording regions. Dark squares indicate statistically significant comparisons (threshold for significance was 0.05/6 for first six columns and 0.05 for final column). All tests were non-parametric (two-sided Wilcoxon rank-sum).

## References

Ahmadlou, M., and Heimel, J.A. (2015). Preference for concentric orientations in the mouse superior colliculus. Nat. Commun. 6, 6773.

Allen, A.E., Procyk, C.A., Howarth, M., Walmsley, L., and Brown, T.M. (2016). Visual input to the mouse lateral posterior and posterior thalamic nuclei: photoreceptive origins and retinotopic order. J. Physiol. 594, 1911–1929.

Baldwin, M.K.L., Wong, P., Reed, J.L., and Kaas, J.H. (2011). Superior colliculus connections with visual thalamus in gray squirrels (Sciurus carolinensis): evidence for four subdivisions within the pulvinar complex. J. Comp. Neurol. 519, 1071–1094.

Baldwin, M.K.L., Balaram, P., and Kaas, J.H. (2013). Projections of the superior colliculus to the pulvinar in prosimian galagos (Otolemur garnettii) and VGLUT2 staining of the visual pulvinar. J. Comp. Neurol. 521, 1664–1682.

Baldwin, M.K.L., Balaram, P., and Kaas, J.H. (2017). The evolution and functions of nuclei of the visual pulvinar in primates. J. Comp. Neurol. 525, 3207–3226.

Bender, D.B. (1983). Visual activation of neurons in the primate pulvinar depends on cortex but not colliculus. Brain Res. 279, 258–261.

Berman, R.A., and Wurtz, R.H. (2010). Functional Identification of a Pulvinar Path from Superior Colliculus to Cortical Area MT. J. Neurosci. 30, 6342–6354.

Berman, R.A., and Wurtz, R.H. (2011). Signals Conveyed in the Pulvinar Pathway from Superior Colliculus to Cortical Area MT. J. Neurosci. 31, 373–384.

Berman, R.A., Cavanaugh, J., McAlonan, K., and Wurtz, R.H. (2017). A circuit for saccadic suppression in the primate brain. J. Neurophysiol. 117, 1720–1735.

Butler, A.B. (2008). Evolution of the thalamus: a morphological and functional review. Thalamus Relat. Syst. 4, 35–58.

Casanova, C., and Molotchnikoff, S. (1990). Influence of the superior colliculus on visual responses of cells in the rabbit’s lateral posterior nucleus. Exp. Brain Res. 80, 387–396.

Chomsung, R.D., Petry, H.M., and Bickford, M.E. (2008). Ultrastructural examination of diffuse and specific tectopulvinar projections in the tree shrew. J. Comp. Neurol. 510, 24–46.

Collins, D.P., Anastasiades, P.G., Marlin, J.J., and Carter, A.G. (2018). Reciprocal Circuits Linking the Prefrontal Cortex with Dorsal and Ventral Thalamic Nuclei. Neuron 98, 366–379.e4.

de Vries, S.E.J., and Clandinin, T.R. (2012). Loom-Sensitive Neurons Link Computation to Action in the Drosophila Visual System. Curr. Biol. 22, 353–362.

Drager, U.C., and Hubel, D.H. (1975). Responses to visual stimulation and relationship between visual, auditory, and somatosensory inputs in mouse superior colliculus. J. Neurophysiol. 38, 690–713.

Dräger, U.C., and Hubel, D.H. (1976). Topography of visual and somatosensory projections to mouse superior colliculus. J. Neurophysiol. 39, 91–101.

Durand, S., Iyer, R., Mizuseki, K., de Vries, S., Mihalas, S., and Reid, R.C. (2016). A Comparison of Visual Response Properties in the Lateral Geniculate Nucleus and Primary Visual Cortex of Awake and Anesthetized Mice. J. Neurosci. 36, 12144–12156.

Frost, B.J., Cavanagh, P., and Morgan, B. (1988). Deep tectal cells in pigeons respond to kinematograms. J. Comp. Physiol. A. 162, 639–647.

Gabbiani, F., Krapp, H.G., and Laurent, G. (1999). Computation of object approach by a wide-field, motion-sensitive neuron. J. Neurosci. 19, 1122–1141.

Gale, S.D., and Murphy, G.J. (2014). Distinct Representation and Distribution of Visual Information by Specific Cell Types in Mouse Superficial Superior Colliculus. J. Neurosci. 34, 13458–13471.

Gale, S.D., and Murphy, G.J. (2016). Active Dendritic Properties and Local Inhibitory Input Enable Selectivity for Object Motion in Mouse Superior Colliculus Neurons. J. Neurosci. 36, 9111–9123.

Gale, S.D., and Murphy, G.J. (2018). Distinct cell types in the superficial superior colliculus project to the dorsal lateral geniculate and lateral posterior thalamic nuclei. J. Neurophysiol. jn.00248.2018.

Garrett, M.E., Nauhaus, I., Marshel, J.H., and Callaway, E.M. (2014). Topography and Areal Organization of Mouse Visual Cortex. J. Neurosci. 34, 12587–12600.

Gerfen, C.R., Paletzki, R., and Heintz, N. (2013). GENSAT BAC Cre-Recombinase Driver Lines to Study the Functional Organization of Cerebral Cortical and Basal Ganglia Circuits. Neuron 80, 1368–1383.

Groh, A., Bokor, H., Mease, R.A., Plattner, V.M., Hangya, B., Stroh, A., Deschenes, M., and Acsády, L. (2014). Convergence of Cortical and Sensory Driver Inputs on Single Thalamocortical Cells. Cereb. Cortex 24, 3167–3179.

Guo, Z. V., Inagaki, H.K., Daie, K., Druckmann, S., Gerfen, C.R., and Svoboda, K. (2017). Maintenance of persistent activity in a frontal thalamocortical loop. Nature 545, 181–186.

Guo, Z.V., Li, N., Huber, D., Ophir, E., Gutnisky, D., Ting, J.T., Feng, G., and Svoboda, K. (2014). Flow of Cortical Activity Underlying a Tactile Decision in Mice. Neuron 81, 179–194.

Halassa, M.M., and Kastner, S. (2017). Thalamic functions in distributed cognitive control. Nat. Neurosci. 20, 1669–1679.

Harris, J.A., Mihalas, S., Hirokawa, K.E., Whitesell, J.D., Knox, J., Bernard, A., Bohn, P., Caldejon, S., Casal, L., Cho, A., et al. (2018). The organization of intracortical connections by layer and cell class in the mouse brain. BioRxiv 292961.

Jun, J.J., Steinmetz, N.A., Siegle, J.H., Denman, D.J., Bauza, M., Barbarits, B., Lee, A.K., Anastassiou, C.A., Andrei, A., Aydın, Ç., et al. (2017). Fully integrated silicon probes for high-density recording of neural activity. Nature 551, 232–236.

Kaas, J.H., and Lyon, D.C. (2007). Pulvinar contributions to the dorsal and ventral streams of visual processing in primates. Brain Res. Rev. 55, 285–296.

Liu, Y.-J., Wang, Q., and Li, B. (2011). Neuronal Responses to Looming Objects in the Superior Colliculus of the Cat. Brain. Behav. Evol. 77, 193–205.

Lyon, D.C., Nassi, J.J., and Callaway, E.M. (2010). A disynaptic relay from superior colliculus to dorsal stream visual cortex in macaque monkey. Neuron 65, 270–279.

Madisen, L., Mao, T., Koch, H., Zhuo, J., Berenyi, A., Fujisawa, S., Hsu, Y.-W.A., Garcia, A.J., Gu, X., Zanella, S., et al. (2012). A toolbox of Cre-dependent optogenetic transgenic mice for light-induced activation and silencing. Nat. Neurosci. 15, 793–802.

Mease, R.A., Sumser, A., Sakmann, B., and Groh, A. (2016). Cortical Dependence of Whisker Responses in Posterior Medial Thalamus In Vivo. Cereb. Cortex 26, 3534–3543.

Moran, P.A.P. (1950). Notes on Continuous Stochastic Phenomena. Biometrika 37, 17.

Nakamura, H., Hioki, H., Furuta, T., and Kaneko, T. (2015). Different cortical projections from three subdivisions of the rat lateral posterior thalamic nucleus: A single-neuron tracing study with viral vectors. Eur. J. Neurosci. 41, 1294–1310.

Oh, S.W., Harris, J.A., Ng, L., Winslow, B., Cain, N., Mihalas, S., Wang, Q., Lau, C., Kuan, L., Henry, A.M., et al. (2014). A mesoscale connectome of the mouse brain. Nature 508, 207–214.

Pachitariu, M., Steinmetz, N.A., Kadir, S.N., Carandini, M., and Harris, K.D. (2016). Fast and accurate spike sorting of high-channel count probes with KiloSort. Adv. Neural Inf. Process. Syst. 29, 4448–4456.

Petersen, S.E., Robinson, D.L., and Keys, W. (1985). Pulvinar nuclei of the behaving rhesus monkey: visual responses and their modulation. J. Neurophysiol. 54, 867–886.

Purushothaman, G., Marion, R., Li, K., and Casagrande, V.A. (2012). Gating and control of primary visual cortex by pulvinar. Nat. Neurosci. 15, 905–912.

Rossant, C., Kadir, S.N., Goodman, D.F.M., Schulman, J., Hunter, M.L.D., Saleem, A.B., Grosmark, A., Belluscio, M., Denfield, G.H., Ecker, A.S., et al. (2016). Spike sorting for large, dense electrode arrays. Nat. Neurosci. 19, 634–641.

Roth, M.M., Dahmen, J.C., Muir, D.R., Imhof, F., Martini, F.J., and Hofer, S.B. (2016). Thalamic nuclei convey diverse contextual information to layer 1 of visual cortex. Nat. Neurosci. 19, 299–307.

Saalmann, Y.B., Pinsk, M.A., Wang, L., Li, X., and Kastner, S. (2012). The Pulvinar Regulates Information Transmission Between Cortical Areas Based on Attention Demands. Science (80-.). 337, 753–756.

Schmitt, L.I., Wimmer, R.D., Nakajima, M., Happ, M., Mofakham, S., and Halassa, M.M. (2017). Thalamic amplification of cortical connectivity sustains attentional control. Nature 545, 219–223.

Shang, C., Chen, Z., Liu, A., Li, Y., Zhang, J., Qu, B., Yan, F., Zhang, Y., Liu, W., Liu, Z., et al. (2018). Divergent midbrain circuits orchestrate escape and freezing responses to looming stimuli in mice. Nat. Commun. 9, 1232.

Sherman, S.M. (2016). Thalamus plays a central role in ongoing cortical functioning. Nat. Neurosci. 16, 533–541.

Shipp, S. (2003). The functional logic of cortico-pulvinar connections. Philos. Trans. R. Soc. Lond. B. Biol. Sci. 358, 1605–1624.

Siegle, J.H., López, A.C., Patel, Y.A., Abramov, K., Ohayon, S., and Voigts, J. (2017). Open Ephys: an open-source, plugin-based platform for multichannel electrophysiology. J. Neural Eng. 14, 045003.

Sun, H., and Frost, B.J. (1998). Computation of different optical variables of looming objects in pigeon nucleus rotundus neurons. Nat. Neurosci. 1, 296–303.

Takahashi, T. (1985). The organization of the lateral thalamus of the hooded rat. J. Comp. Neurol. 231, 281–309.

Tohmi, M., Meguro, R., Tsukano, H., Hishida, R., and Shibuki, K. (2014). The extrageniculate visual pathway generates distinct response properties in the higher visual areas of mice. Curr. Biol. 24, 587–597.

Wang, Q., and Burkhalter, A. (2013). Stream-Related Preferences of Inputs to the Superior Colliculus from Areas of Dorsal and Ventral Streams of Mouse Visual Cortex. J. Neurosci. 33, 1696–1705.

Wang, Q., Sporns, O., and Burkhalter, A. (2012). Network Analysis of Corticocortical Connections Reveals Ventral and Dorsal Processing Streams in Mouse Visual Cortex. J. Neurosci. 32, 4386–4399.

Wei, P., Liu, N., Zhang, Z., Liu, X., Tang, Y., He, X., Wu, B., Zhou, Z., Liu, Y., Li, J., et al. (2015). Processing of visually evoked innate fear by a non-canonical thalamic pathway. Nat. Commun. 6, 6756.

Wickersham, I.R., Lyon, D.C., Barnard, R.J.O., Mori, T., Finke, S., Conzelmann, K.-K., Young, J.A.T., and Callaway, E.M. (2007). Monosynaptic Restriction of Transsynaptic Tracing from Single, Genetically Targeted Neurons. Neuron 53, 639–647.

Zhao, S., Ting, J.T., Atallah, H.E., Qiu, L., Tan, J., Gloss, B., Augustine, G.J., Deisseroth, K., Luo, M., Graybiel, A.M., et al. (2011). Cell type–specific channelrhodopsin-2 transgenic mice for optogenetic dissection of neural circuitry function. Nat. Methods 8, 745–752.

Zhao, X., Liu, M., and Cang, J. (2014). Visual Cortex Modulates the Magnitude but Not the Selectivity of Looming-Evoked Responses in the Superior Colliculus of Awake Mice. Neuron 84, 202–213.

Zhou, H., Schafer, R.J., and Desimone, R. (2016). Pulvinar-Cortex Interactions in Vision and Attention. Neuron 89, 209–220.

Zhou, N., Maire, P.S., Masterson, S.P., and Bickford, M.E. (2017). The mouse pulvinar nucleus: Organization of the tectorecipient zones. Vis. Neurosci. 34, E011.

Zhuang, J., Ng, L., Williams, D., Valley, M., Li, Y., Garrett, M., and Waters, J. (2017). An extended retinotopic map of mouse cortex. Elife 6.

